# Disrupted chromatin architecture blocks B lymphopoiesis in Down syndrome

**DOI:** 10.64898/2026.10.07.757425

**Authors:** Zhicong Liao, Mimi Zhang, Brian Y. Soong, Bohan Zhu, Linda L. Boshans, Ke Wang, Miguel Quijada-Álamo, Ifeoma Osakwe, Zhe Wang, Persephone Fiore, Dan Hasson, Elvin Wagenblast, Sai Ma

## Abstract

Human fetal haematopoiesis requires developmental potential to be encoded before lineage identity becomes transcriptionally apparent, but whether three-dimensional genome architecture provides such information is unknown. Here we develop ChromE-seq, a scalable single-cell assay that jointly profiles chromatin conformation and transcription, and integrate it with chromatin accessibility and DNA methylation to generate a multimodal atlas of ∼400,000 haematopoietic cells from disomic and trisomy 21 (T21) human fetal liver. Chromatin contacts frequently preceded lineage-specific transcription or persisted after transcription declined, revealing 3D chromatin priming and memory. Unexpectedly, B-lineage and erythroid differentiation followed opposing architectural trajectories, with progressive loss and gain of chromatin connectivity, respectively. T21 disproportionately disrupted B-lineage architecture, redistributed interactions towards shorter genomic distances and weakened an EBF1-centred regulatory circuit. Deletion of a T21-sensitive EBF1 loop anchor reproduced the CLP-to-B-progenitor bottleneck observed in T21. These findings establish 3D genome architecture as an organizing layer of human fetal haematopoiesis that links developmental potential with lineage-specific vulnerability to genomic perturbation.

**Highlights:**

- A ∼400,000-cell multimodal atlas resolves human fetal haematopoietic genome regulation
- 3D chromatin encodes developmental priming and persistence
- Trisomy 21 selectively disrupts B-lineage genome architecture
- A T21-sensitive *EBF1* loop is required for normal B-cell development

## Main

Human fetal haematopoiesis requires multipotent stem cells and progenitors to generate distinct blood and immune lineages while progressively restricting developmental potential. Transcriptional state alone may not fully capture this potential, because epigenetic configurations can be established long before gene activation or persist after transcription has declined^1^. Chromatin accessibility can anticipate gene activation during haematopoietic differentiation^1–3^, but it is unknown whether three-dimensional (3D) genome architecture carries comparable temporal information. By bringing distal regulatory elements into physical proximity with their target genes, chromatin folding could encode future developmental potential and preserve features of earlier cell states. How chromatin contacts are temporally coordinated with transcription, and whether this relationship differs across human fetal blood lineages, remains unresolved.

B lymphopoiesis provides a particularly informative setting in which to address this question. Distal regulatory interactions contribute to B-lineage specification^4^, raising the possibility that B-cell differentiation may depend on a distinctive programme of 3D genome reorganization. This question is especially relevant to trisomy 21 (T21), known as Down syndrome, a constitutional aneuploidy present in every cell yet with strikingly lineage-selective consequences in haematopoiesis. T21 alters erythroid and megakaryocytic development and dramatically impairs B lymphopoiesis^1,5–7^, while strongly predisposing to childhood leukaemia^8–10^. This B-cell defect is already evident before birth, with severe depletion of committed early B-cell progenitors despite relative preservation of upstream progenitors^5,6^. Strikingly, this developmental block coexists with an approximately 20-fold increased risk of acute lymphoblastic leukaemia (ALL), which in Down syndrome is almost exclusively B-lineage^11^. What makes the developing B-lineage particularly vulnerable to chromosome 21 dosage? We reasoned that the answer might lie in the 3D genome architecture established during fetal haematopoiesis.

Here we develop **Chrom**atin and **E**xpression co-assay with **seq**uencing (ChromE-seq), a scalable single-cell assay that jointly profiles chromatin conformation and transcription, and integrate it with chromatin accessibility, DNA methylation, and spatial transcriptomics across ∼400,000 haematopoietic cells from disomic and T21 human fetal liver. We uncover 3D chromatin-associated priming and memory, in which chromatin contacts anticipate lineage-specific transcription or persist after transcriptional programs decline. Unexpectedly, B-lineage and erythroid differentiation adopt opposing architectural trajectories, characterized by progressive loss and gain of chromatin connectivity, respectively. T21 disproportionately disrupts B-lineage architecture, redistributing chromatin interactions toward shorter genomic distances and weakening an EBF1-centred regulatory circuit. Deletion of a T21-sensitive *EBF1* loop anchor reproduces a common lymphoid progenitor (CLP)-to-B-progenitor bottleneck observed in T21. Together, these findings establish 3D genome architecture as an organizing layer of human fetal haematopoiesis that encodes developmental competence and shapes lineage-specific vulnerability to genomic perturbation.

### ChromE-seq jointly profiles genome architecture and transcription at scale

We developed ChromE-seq, a scalable single-cell assay that combines *in situ* restriction digestion and proximity ligation with reverse transcription and combinatorial indexing to jointly measure chromatin contacts and RNA in the same cells (**Extended Data Fig. 1**). After extensive optimization for cellular recovery and molecular yield (**Extended Data Fig. 2a–e, Extended Data Table 1**), human-mouse mixing experiments showed concordant species assignment from both modalities, supporting the specificity of cellular barcoding (**Extended Data Fig. 1b, c, Table 2**). Paired profiles further recovered concordant cell populations in mouse brain, and aggregated fetal haematopoietic stem and progenitor cell (HSPC) contact maps recapitulated major features of published bulk Hi-C data (**Extended Data Figs. 1d and 2f–h**). Across benchmarked conditions, ChromE-seq achieved high cellular throughput with chromatin contact and transcript recovery comparable to existing joint conformation-transcription assays^12–16^, at an estimated ∼17-fold lower cost per cell than the lowest-cost existing single-cell Hi-C assay (**Extended Data Table 3, 4, Extended Data Fig. 1f–i**). Together, these benchmarks established ChromE-seq as a scalable platform for paired single-cell profiling of chromatin conformation and transcription. We next applied it to human fetal liver haematopoiesis, where rare stem, progenitor, and transitional states can only be resolved through deep cellular sampling across multiple donors.

### A conformation-resolved atlas of fetal haematopoiesis

To dissect how chromosomal conformation affects early blood development, we profiled CD34^+^ HSPCs together with CD45^+^ immune cells and CD45^−^ stromal cells from seven disomic (D21) and five T21 human fetal liver specimens using ChromE-seq and ME-seq^17^, generating 121,367 paired Hi-C–RNA and 278,401 paired ATAC–methylation–RNA single-cell profiles, representing the largest single-cell atlas incorporating paired chromatin-conformation and transcription measurements in haematopoiesis to date, which we refer to as **Hu**man **F**etal **H**aematopoiesis **A**tlas (HuFHA, **Fig. 1a, b, Extended Data Fig. 3a, Extended Data Table 5**). To provide tissue context, we also performed Visium HD spatial transcriptomics on matched fetal liver sections.

**Figure 1.**
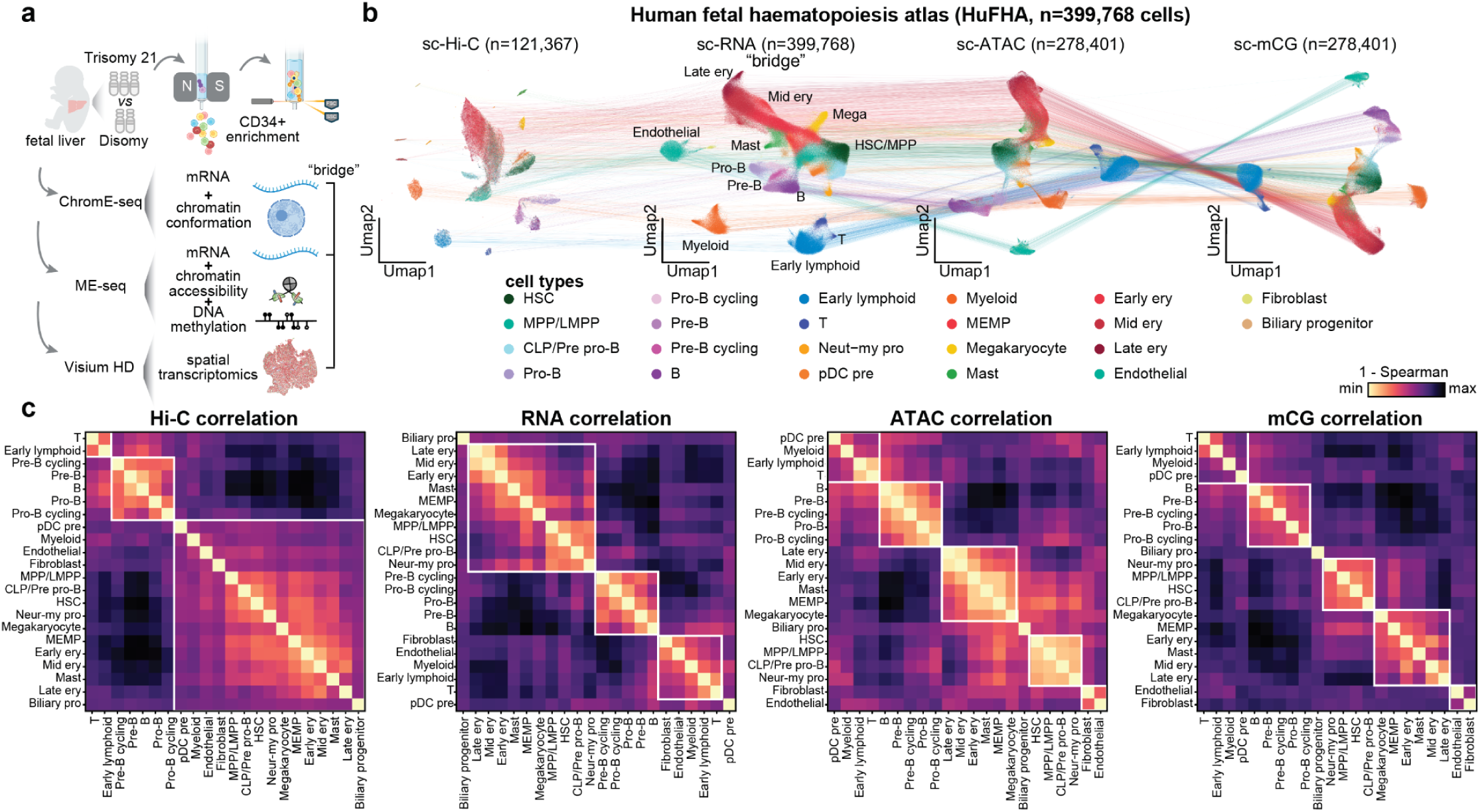
A multi-modal single-cell atlas of human fetal haematopoiesis. **a**, Schematic of study design to comprehensively profile HSPCs, immune cells and stromal cells from disomy and trisomy 21 fetal livers using ChromE-seq and ME-seq. **b**, Integrated UMAP for the four single-cell modalities, with cell types coloured based on RNA-derived annotations. Lines indicate modalities measured in the same cell. **c**, Pairwise Spearman correlation between cell types calculated separately for each modality. White boxes indicate major lineage-associated groups.

Integration through shared RNA modality resolved 22 cell types, including 19 haematopoietic populations (**Fig. 1b**). Cell identities were supported by canonical lineage markers (**Extended Data Fig. 4**) and validated against published fetal and adult human haematopoietic references (**Extended Data Fig. 3b–d**). Chromosome 21 dosage was independently recovered from both RNA- and Hi-C-derived copy-number profiles, and multimodal differential analysis in haematopoietic stem cells (HSC)/multipotent progenitor cells (MPP) revealed coordinated T21-associated changes across transcription, chromatin accessibility, DNA methylation, and chromatin contacts (**Extended Data Fig. 5**). Disomic and T21 donors were well represented across the integrated atlas, with comparable per-donor data quality across modalities (**Extended Data Fig. 5b–d**). As expected, across modalities, lineage progression was accompanied by progressive activation of lineage-associated transcription, increased chromatin accessibility, reduced methylation, and reorganization of chromatin contacts at lineage-specific regulatory regions (**Extended Data Fig. 4a, c**). ChromE-seq further resolved cell-type-specific contact patterns at lineage-regulatory loci, including distinct A/B compartment organization and looping configuration around the canonical B-lineage marker gene *EBF1* across haematopoietic populations (**Extended Data Fig. 4b**).

We next asked whether lineage relationships were represented similarly across regulatory layers. Interestingly, RNA expression, chromatin accessibility, and DNA methylation largely preserved the expected haematopoietic hierarchy, whereas chromatin conformation revealed a pronounced separation of B-lineage cells from other populations (**Fig. 1c**). These findings indicate that fetal B-cell differentiation is accompanied by a distinct reorganization of 3D genome architecture, raising the question of how this regulatory transition is established during normal fetal haematopoiesis and is perturbed by genomic abnormality.

### 3D chromatin priming and memory during differentiation

Epigenetic states can precede transcriptional activation or persist after transcriptional programmes decline, reflecting regulatory priming and memory. Prior studies of fetal haematopoiesis have shown that chromatin accessibility can precede transcriptional activation, providing evidence for regulatory priming. Whether 3D chromosome conformation shows similar temporal offsets, including both priming before gene activation and persistence after transcriptional programmes decline, remains unclear. We therefore reconstructed B-lymphoid and erythroid trajectories, enabling comparison of chromatin-contact and transcriptional dynamics along lineage progression (**Fig. 2a**).

**Figure 2.**
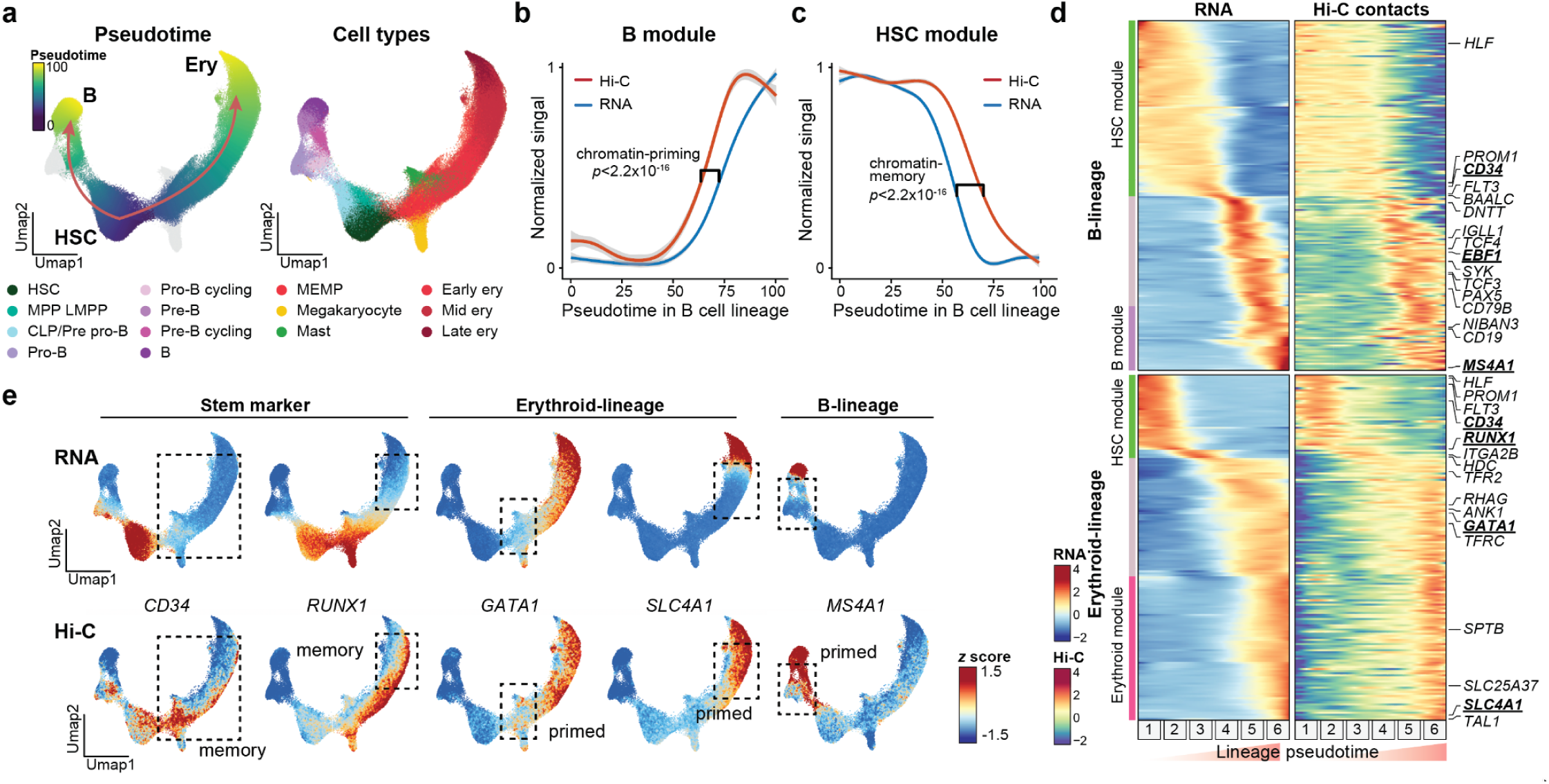
3D chromatin priming and memory along fetal haematopoietic differentiation. **a**, Pseudotime reconstruction of B and erythroid differentiation trajectories, shown by pseudotime and annotated cell types. **b,c,** Normalized Hi-C contacts and gene expression signal along B-lineage pseudotime for a B-lineage-associated module (**b**) and an HSC-associated module (**c**). Lines indicate smoothed trends; shaded regions indicate 95% confidence interval. **d**, Heatmap of normalized Hi-C and gene expression signal over inferred pseudotime for unbiasedly identified gene modules along B and erythroid lineages, with selected lineage marker genes highlighted. **e**, Normalized Hi-C contacts and gene expression signal for representative stem/progenitor (*CD34*, *RUNX1*), erythroid (*GATA1*, *SLC4A1*), and B-lineage (*MS4A1*) genes, which exhibit chromatin priming and memory.

Unsupervised clustering of genes along pseudotime identified lineage-associated gene modules with distinct temporal relationships between chromatin contacts and transcription (**Fig. 2b–d, Extended Data Fig. 6**). Notably, along the B-cell trajectory, lineage-associated chromatin contacts increased significantly prior to the onset of gene expression, consistent with 3D chromatin priming (**Fig. 2b**, *p* < 2.2×10^−16^, *K-S* test). Conversely, contacts associated with the HSC module persisted well past the decline of transcription, revealing chromatin memory during lineage exit (**Fig. 2c**, *p* < 2.2×10^−16^, *K-S* test). Similar but more pronounced temporal offsets were also observed along the erythroid trajectory (**Extended Data Fig. 6f, g**).

These temporal relationships were also evident at lineage-defining loci. Stem-associated genes such as *CD34* and *RUNX1* retained elevated chromatin contact signals after their transcriptional programmes diminished, whereas lineage genes including erythroid genes (*GATA1*, *SLC4A1*) and a mature B-cell gene (*MS4A1*) acquired increased chromatin contacts ahead of transcriptional activation (**Fig. 2e**). Together, our results demonstrate how 3D genome organization can both anticipate future lineage programmes and retain features of earlier cell states.

Given the substantial 3D chromatin changes accompanying B-lineage differentiation, we focused on the B-lineage differentiation regulator *EBF1*. Indeed, this temporal ordering was also evident at the *EBF1* locus, where Hi-C-derived A/B compartments and distal regulatory loops changed before the promoter accessibility increase and robust transcriptional induction, while DNA methylation remained comparatively stable (**Extended Data Fig. 6a–e**). Progressive remodelling of the *EBF1 cis*-regulatory architecture preceded the rise in *EBF1* expression as cells entered the CLP/pre-pro-B state, indicating that the architectural transition is established ahead of B-lineage transcriptional commitment. Together, these data indicate the dependence of B-cells on their chromatin structure for lineage commitment, and underscore the need to measure multiple epigenetic modalities to fully resolve developmental transitions obscured when a single modality is considered alone.

### Trisomy 21 disrupts B-lineage genome architecture

B lymphopoiesis is selectively impaired in Down syndrome^5,6^, and we therefore asked whether the distinctive 3D architecture that accompanies B-cell differentiation contributes to this defect. Consistent with previous studies^7^, B-lineage cells are significantly depleted in T21 fetal liver in our atlas (**Fig. 3a, b**, *p* < 0.01, *t*-test). Flow cytometry analysis of CD45^+^CD34^+^ cells from disomic and T21 fetal liver confirmed this loss, and further revealed a significant accumulation of CLPs alongside a reduction in downstream committed B-cell progenitors, defined as the combined pre-pro-B and pro-B population (**Fig. 3c, Extended Data Fig. 7a**). Immunohistochemical (IHC) staining of T21 fetal liver sections likewise showed reduced CD19^+^ B-cells, confirming the depletion in intact tissue (**Extended Data Fig. 7b, c**). Together, these results localize the bottleneck at the transition from CLP to committed B-cell progenitors and suggest it may be epigenetically imposed.

**Figure 3.**
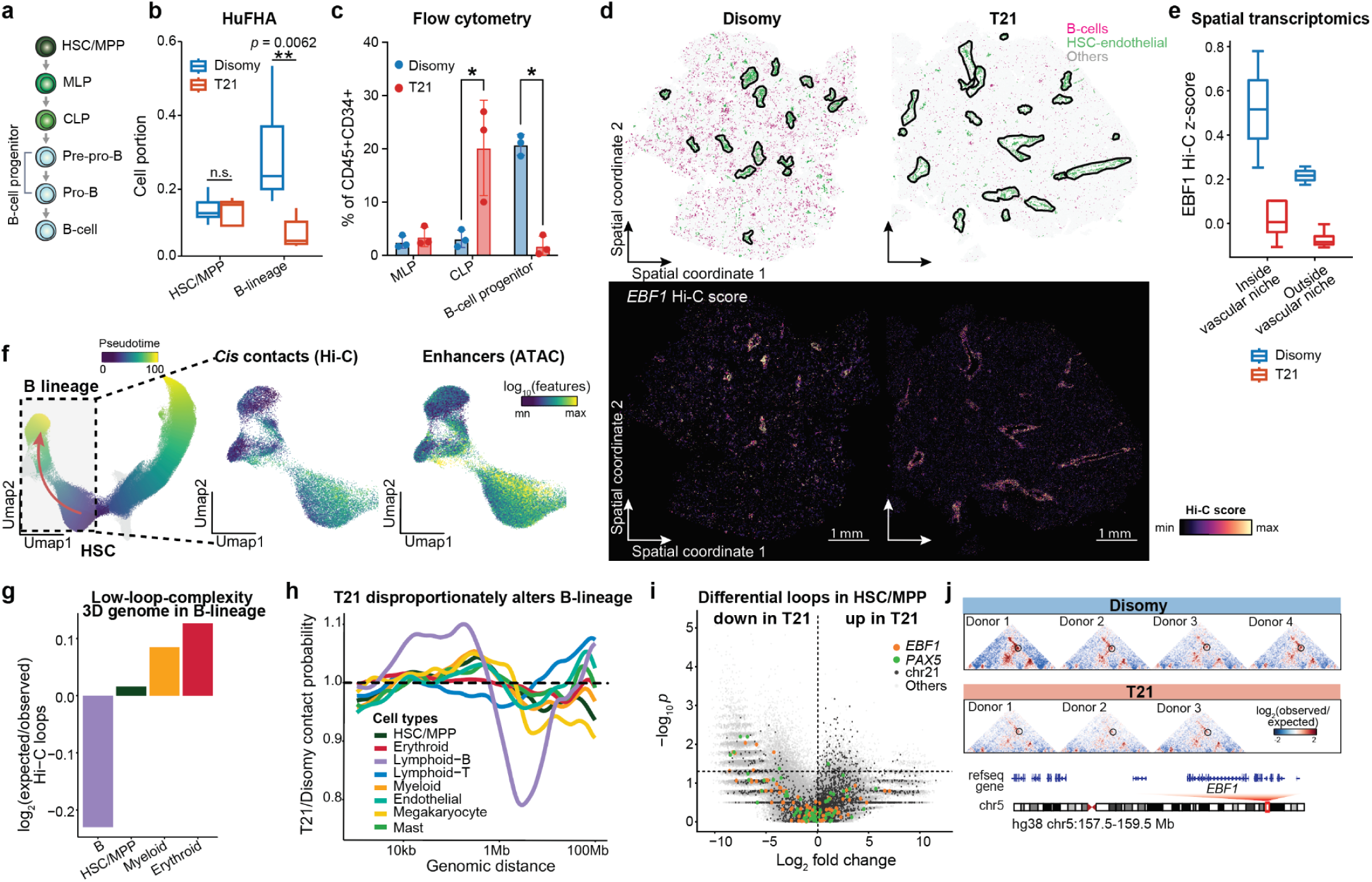
Trisomy 21 selectively disrupts B-lineage genome architecture. **a**, Schematic of B-cell differentiation. The B-cell progenitor population comprises the combined pre-pro-B and pro-B compartments used throughout this study. **b,** Relative abundance of HSC/MPP and B-lineage cells in the single-cell atlas. **c**, Quantification of multi-lymphoid progenitors (MLPs), common lymphoid progenitors (CLPs) and downstream B-cell progenitors among CD45^+^CD34^+^ cells in disomic and T21 fetal liver by flow cytometry. Population definitions and the complete gating strategy are shown in Extended Data Fig. 7a. **d**, Representative Visium HD visualizations for disomic and T21 samples, highlighting HSCs and B-cells, with endothelial-associated HSC niches outlined in black, along with projected Hi-C *EBF1* score. **e**, Projected *EBF1* Hi-C signature in Visium HD spots located within or outside HSC-associated vascular niches in disomic and T21 fetal liver. **f,** Visualization of pseudotime trajectory along B-lineage, alongside the observed number of Hi-C *cis* contacts and accessible putative enhancers. **g,** Log-fold differences between expected and observed Hi-C loops for each lineage after balanced downsampling. **h**, Ratio of T21-to-disomy Hi-C contact probability over genomic distances for each lineage. **i**, Differential chromatin loop between T21 and disomic in HSC/MPP, with highlighted loops related to *EBF1*, *PAX5*, and chromosome 21. Dashed lines indicate the statistical significance and fold-change threshold used to define differential loops. **j,** Donor-level Hi-C contact maps across the *EBF1* locus. The T21-sensitive *EBF1* loop is indicated. * *p* < 0.05, ** *p* < 0.01, *t*-test.

To examine how 3D chromatin states are distributed in their native tissue context, we projected ChromE-seq-derived Hi-C signatures onto gestational-matched Visium HD spatial transcriptomic fetal liver sections. This approach localized B-lineage-associated chromatin states and confirmed reduced B-cell representation in T21 fetal liver (**Fig. 3d**). The projected *EBF1* contact signature was likewise reduced in T21, including within HSC-associated vascular niches (**Fig. 3d, e, Extended Data Fig. 8**). Across both IHC and spatial transcriptomic data, B-cell abundance significantly increased with gestational age in disomic fetal liver, as expected with normal fetal development. This increase was substantially attenuated in T21, indicating a prominent deficient B-lineage development in fetal liver tissue (IHC: *p =* 0.011, Visium HD: *p* = 0.0044, gestational-age adjusted linear regression, **Extended Data Fig. 7c, e**). Analysis of transcriptional potency with CytoTRACE 2^18^ further showed significantly reduced scores in T21 progenitors consistent with an early cell-intrinsic defect, indicating that the B-cell deficit is accompanied by broader changes in the stem and progenitor compartment before differentiated B-cells are lost (**Extended Data Fig. 9a, b**).

We next asked whether B-lineage differentiation is associated with a distinctive architectural state that explains its sensitivity to chromosome 21 dosage. Along B-cell differentiation, both accessible regulatory elements and *cis* chromatin contacts progressively declined, suggesting strong repression and compaction of the genome (**Fig. 3f, Extended Data Fig. 9**). In contrast, erythroid differentiation displayed increasing contact frequencies (**Extended Data Fig. 9c**). After balanced downsampling across lineages to control for differences in cell number, B-lineage cells retained significantly fewer chromatin loops than expected (**Fig. 3g**), revealing a low-complexity 3D genome architecture compared with the loop-rich erythroid lineage. Thus, lineage restriction is accompanied by fundamentally different modes of 3D genome remodelling: B-cell development follows a restrictive mode characterized by progressive loss of chromatin contacts and loops, whereas erythroid differentiation adopts a more constructive, loop-building program.

We therefore asked whether this distinctive B-lineage architecture was preferentially disrupted by additional chromosomal perturbation. At higher levels of genome organization, T21 shortened A/B compartments and increased compartment switching across major cell types, and broadly reduced interchromosomal contacts in HSC/MPP, consistent with reduced interchromosomal mixing, or chromosomal introversion^19^ (**Extended Data Fig. 9g-i**). Within chromosomes, T21 B-lineage cells showed a redistribution of chromatin contacts towards shorter genomic distances, with increased short-range (10 kb–1 Mb) interactions and reduced long-range (>1 Mb) contacts compared with disomic cells (**Fig. 3h**). This shift was substantially less pronounced in the other haematopoietic lineages, indicating that T21 disproportionately perturbs B-lineage architecture. Differential loop analysis in HSC/MPP recovered the expected gain of chromosome 21-associated interactions, reflecting the additional copy. Contacts involving key B-cell regulators, including EBF1 and PAX5, were among those reduced in T21 (**Fig. 3i, Extended Data Table 6**). Reduced *EBF1*-associated contacts were also reproducible across individual T21 donors (**Fig. 3j**).

Together, these findings suggest that the restrictive, low-connectivity 3D genome organization accompanying B-cell differentiation is associated with heightened architectural vulnerability to chromosome 21 dosage. In T21, this vulnerability is associated with altered higher-order genome organization, redistribution of chromatin contacts, and loss of regulatory interactions at key B-cell loci, coincident with the developmental bottleneck at CLP-to-B-cell transition. These data motivated us to identify the regulatory interactions most affected by this perturbation.

### T21 rewires B-lineage regulatory priming in stem and progenitors

To define the regulatory circuitry disrupted in T21, we extended our previously described domains of regulatory chromatin (DORC) framework^3^ by integrating accessibility, DNA methylation, transcription, and chromatin conformation. High-resolution peak–gene associations from accessibility–RNA and methylation–RNA coupling were intersected with high-fidelity enhancer–promoter (E–P) loops, which provide direct physical support for candidate distal target assignment, to define loop-assisted DORCs (LA-DORCs; **Fig. 4a**, **Extended Data Fig. 10a**, Methods). Peak–gene associations extended broadly across ±2 Mb of transcription start sites (TSSs), whereas E–P loops identified a more spatially constrained subset of physically supported distal interactions, motivating their integration to refine distal regulatory assignments (**Extended Data Fig. 10b**).

**Figure 4.**
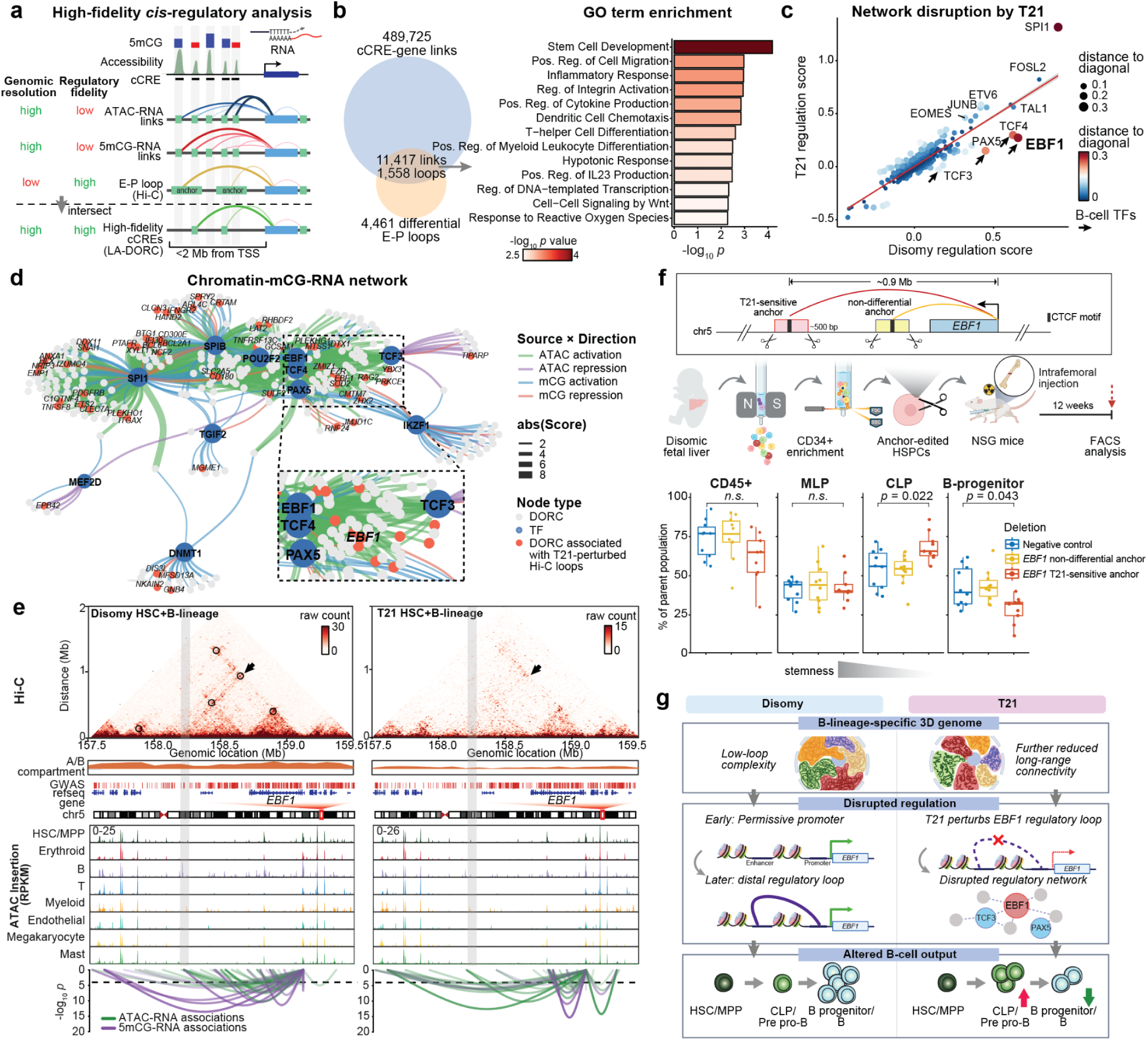
T21 disrupts an EBF1-centred B-lineage regulatory network. **a,** Schematic of loop-assisted DORC (LA-DORC) analysis. High-resolution ATAC–RNA and mCG–RNA peak–gene associations were intersected with enhancer–promoter Hi-C loops to identify distal *cis*-regulatory links supported by physical chromatin contact. **b,** Gene Ontology enrichment of genes associated with T21-differential enhancer–promoter loops intersecting LA-DORC regulatory links. **c,** Comparison of transcription-factor regulation scores between disomic and T21 cells. Point size and colour indicate deviation from the diagonal. **d,** Multimodal regulatory network integrating physical chromatin contacts with accessibility, DNA methylation, transcription-factor motifs and RNA expression. Edge colour denotes regulatory modality and direction, and edge width represents regulation score. Red nodes indicate DORCs associated with T21-perturbed Hi-C loops. Inset highlights the EBF1-centred B-lineage module. **e,** Multimodal regulatory architecture across the *EBF1* locus in disomic and T21 HSC/B-lineage cells. Hi-C contact maps are shown together with compartment state, GWAS annotations, gene models, cell-type-specific chromatin accessibility and ATAC–RNA and mCG–RNA peak–gene associations. Arrows indicate the T21-sensitive *EBF1* regulatory interaction. **f,** Deletion of *EBF1* loop anchors in disomic human fetal liver HSPCs. CD34^+^ HSPCs were edited by CRISPR/Cas9 to delete the T21-sensitive anchor, a T21-insensitive anchor or a control locus, transplanted into NSG mice and analysed by flow cytometry 12 weeks later. Plots show human CD45^+^ cells, MLP, CLP and B-cell progenitors as a percentage of the parent population. Statistical significance was assessed using a two-sided Wilcoxon rank-sum test. **g,** Proposed model in which the relatively low-loop-connectivity B-lineage genome is further disrupted by T21, weakening an EBF1-centred regulatory circuit and producing an early B-lineage differentiation bottleneck.

Among 4,461 HSC/MPP-associated differential E–P loops, 1,558 intersected high-confidence *cis*-regulatory links (**Fig. 4b**). Genes associated with these T21-sensitive interactions were enriched for pathways related to stem cell development, cell migration, inflammatory responses, and lymphocyte differentiation (**Fig. 4b**), indicating that altered chromosome 21 dosage broadly perturbs regulatory programs within early stem and progenitor cells.

To place these findings in a broader regulatory context, we next developed a multimodal regulatory network framework that integrates physical chromatin contacts with accessibility, DNA methylation, transcription factor (TF) expression, and motif occurrence to infer TF–target gene relationships (**Fig. 4c, d**, Methods). Notably, within this network, the *EBF1* LA-DORC was jointly connected to several central B-lineage regulators, including EBF1, PAX5, and TCF-family factors, suggesting that EBF1 sits within a broader B-cell regulatory circuit rather than acting as an isolated target. For each TF, we quantified a regulatory influence score by combining motif enrichment with the association between TF expression and regulatory-element activity, enabling direct comparison between disomic and T21 cells (**Fig. 4c**). Most TFs retained similar regulatory influence in disomic and T21 cells, whereas several core B-lineage regulators, including EBF1, PAX5 and TCF-family factors^4^, showed noticeable loss of regulatory scores in T21. By contrast, a smaller set of myeloid-and activation-associated factors, including SPI1 (PU.1), gained regulatory influence, consistent with T21-associated myeloid bias^20^ and selective network rewiring rather than global loss of regulatory activity (**Fig. 4c**). At the *EBF1* locus itself, the coupling between regulatory-element activity and transcription was also diminished in T21, further prioritizing EBF1 as a candidate mediator of the B-lineage defect. Consistent with network-level disruption, EBF1 showed among the strongest coupling between TF activity and transcription in disomic HSPCs, whereas this relationship was attenuated in T21 (**Extended Data Fig. 10c**). Together, these analyses show that T21 selectively weakens an EBF1-centred B-cell regulatory network and nominates EBF1-associated chromatin regulation for functional investigation.

### Disruption of a T21-sensitive EBF1 regulatory loop impairs B-cell differentiation

Across B-lineage pseudotime, the *EBF1* promoter remained largely unmethylated and accessible before robust transcriptional induction, whereas distal regulatory elements and chromatin contacts were progressively remodelled as cells entered the CLP/pre-pro-B state (**Extended Data Fig. 6a–e**). This temporal ordering suggests that promoter accessibility and methylation alone do not fully account for the onset of *EBF1* expression and that distal chromatin remodelling may play a role in establishing the EBF1-associated regulatory program.

At the *EBF1* locus, T21 selectively weakened a distal E–P interaction (∼0.9 Mb from the *EBF1* promoter, **Fig. 4e**, grey bar) that was prominent in disomic B-lineage cells across donors (**Fig. 3i**), consistent with the broader loss of contact–transcription coupling observed for *EBF1.* Because this regulatory element lies nearly 1 Mb from *EBF1*, its connection to the gene is difficult to infer from chromatin accessibility alone, underscoring the value of directly measuring long-range chromatin contacts. The same region also showed reduced peak–gene association with *EBF1* in T21, providing convergent evidence for disruption of this *cis*-regulatory connection. We therefore focused on this T21-sensitive loop anchor as a candidate regulatory element important for normal B-cell development.

To test its functional contribution directly, we deleted the *EBF1* loop-anchor region, which contains a CTCF motif, in CD34^+^ HSPCs from human fetal liver and assessed lineage output following xenotransplantation in NSG mice (**Fig. 4f**). In parallel, we included the olfactory receptor OR2W5 as a negative control and deleted a second *EBF1* E–P loop anchor that was not differentially altered in T21 (**Extended Data Fig. 11a)**. PCR genotyping of the transplanted cells confirmed the expected anchor deletions, with representative ICE analyses showing high editing efficiencies at both loci (96% and 100% indel frequencies for the T21-sensitive and non-differential anchors, respectively^21^; **Extended Data Fig. 11b–d**). Deletion of the T21-sensitive anchor largely preserved human engraftment (*p* = 0.11, Wilcoxon rank-sum test) and early CD34^+^ HSPC and multi-lymphoid progenitor (MLP) compartments, but altered subsequent B-cell progression. CLPs accumulated, whereas downstream committed B-cell progenitors were reduced, recapitulating the CLP-to-B-lineage bottleneck observed in T21 fetal liver. This phenotype was not observed following deletion of the control locus or the T21-insensitive *EBF1* loop anchor, demonstrating that it reflects loss of this specific regulatory interaction rather than a nonspecific perturbation at the *EBF1* locus. Therefore, deletion of a single architectural element is sufficient to reproduce a central feature of the T21 B-lineage defect in human primary cells.

Together, our multimodal atlas and functional perturbation experiments support a model (**Fig. 4g**) in which fetal B-lineage differentiation is accompanied by progressive restriction of 3D chromatin contacts, elevating sensitivity to disruption of specific regulatory interactions. T21 further reduces long-range connectivity and disrupts specific regulatory interactions at key B-lineage loci, including a distal EBF1 loop that is essential for efficient progression beyond the CLP stage. At the *EBF1* locus, the promoter is permissive before induction, and remodelling of distal contacts precedes robust transcription, activating an EBF1-centred regulatory network involving TCF3/4 and PAX5. Disruption of this network in T21 results in accumulation of CLPs and reduced progression into downstream B-progenitors and mature B-cells, linking altered 3D genome organization to the developmental bottleneck observed in T21.

## Discussion

ChromE-seq, integrated with ME-seq and spatial transcriptomics, enabled us to connect 3D genome organization with accessibility, DNA methylation, transcription, and tissue context across ∼400,000 haematopoietic cells from human fetal liver. This multimodal framework revealed that genome architecture provides a temporal and lineage-specific regulatory layer that is not captured by transcription alone. Chromatin contacts frequently preceded lineage-specific transcription or persisted after transcriptional programs declined, extending the concept of developmental priming beyond local chromatin state. These temporal offsets suggest that genome architecture can encode both future and previous regulatory states, with developmental potential emerging from asynchronous remodelling across chromatin organization, accessibility, DNA methylation, and transcription.

Unexpectedly, lineage commitment followed distinct architectural trajectories. B-lineage differentiation progressively lost *cis* contacts and chromatin loops, whereas erythroid differentiation gained connectivity, revealing opposing modes of genome remodelling during lineage restriction. The resulting low-connectivity B-lineage state may carry less architectural redundancy, such that individual regulatory interactions assume greater importance as differentiation proceeds. This provides a framework for understanding why the same genomic perturbation can have unequal consequences across developing lineages.

T21 exposed this architectural vulnerability. Although chromosome 21 dosage is altered across haematopoietic cells, T21 disproportionately remodelled the B-lineage genome, redistributing contacts towards shorter genomic distances and weakening interactions at key lineage regulators including *EBF1* and *PAX5*. Deletion of a T21-sensitive *EBF1* loop anchor reproduced the CLP-to-B-progenitor bottleneck *in vivo*, whereas deletion of the T21-insensitive anchor did not. In these xenografts, CD19^+^ B-cells were only modestly affected, indicating that deletion of this element slows progression through the CLP transition rather than depleting the mature B-cell compartment. These findings link a specific architecture-defined regulatory element to the developmental phenotype and indicate that 3D genome organization can contribute directly to lineage progression.

Broadly, our findings suggest that the consequences of genomic perturbation depend not only on the altered chromosome itself, but also on the developmental architecture onto which that perturbation is imposed. In T21, this interaction is already evident during fetal haematopoiesis, placing the origin of the B-cell defect before birth and connecting a lifelong immune phenotype to altered developmental genome organization. Whether the same constrained, *EBF1*-weakened architecture also provides a permissive substrate for the B-lineage leukaemic transformation that arises at increased frequency in Down syndrome remains to be tested. Similar principles may help explain selective cellular vulnerability in other aneuploidies, copy-number disorders, and developmental diseases.

## Extended data figures

**Extended Data Figure 1.**
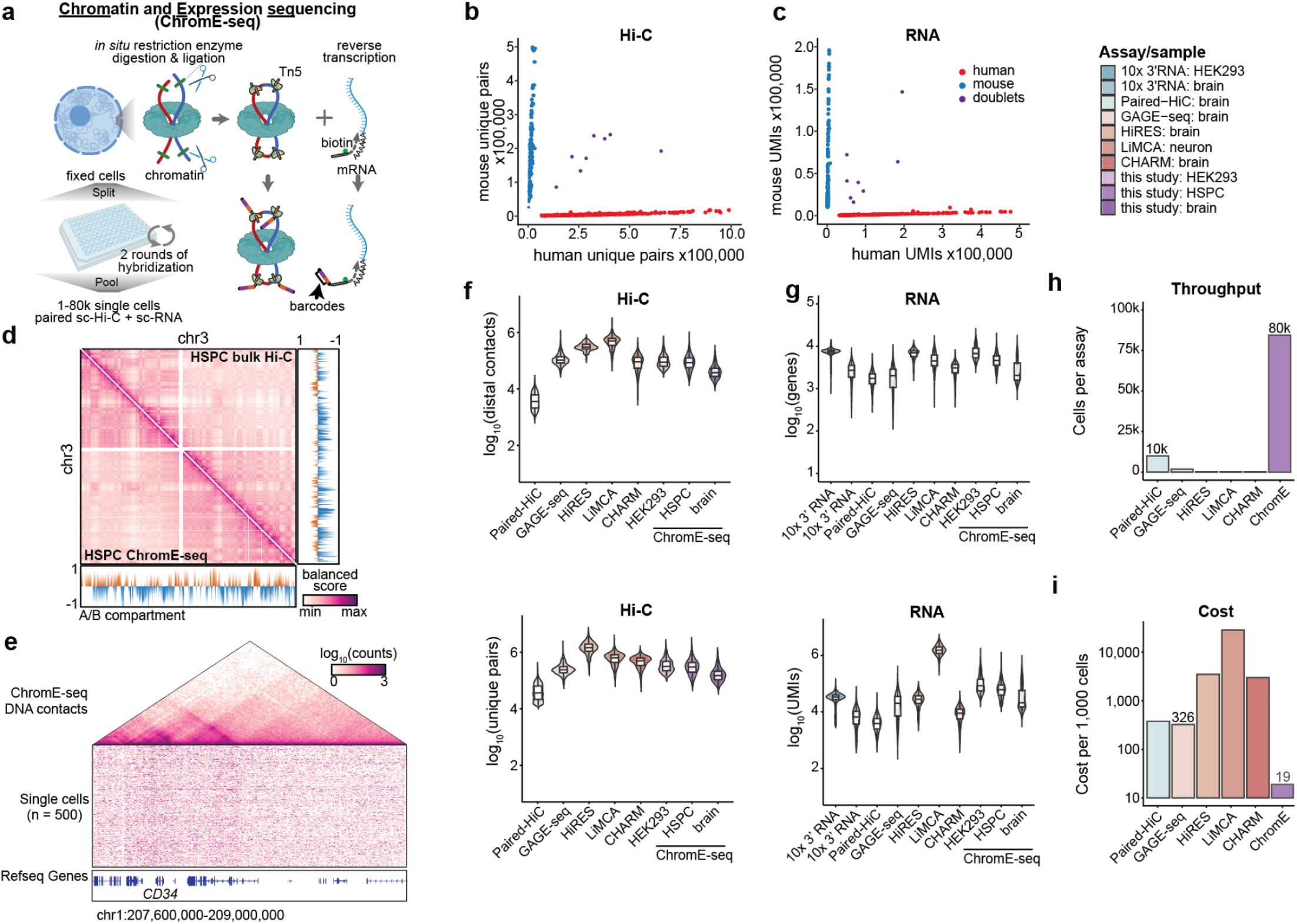
Development and benchmarking of ChromE-seq. **a,** Schematic of the ChromE-seq workflow. Fixed cells undergo *in situ* restriction digestion and proximity ligation for chromatin-contact capture, together with reverse transcription of mRNA, followed by combinatorial indexing to generate paired single-cell Hi-C and RNA profiles. **b,c,** Human–mouse mixing experiment showing species assignment from Hi-C unique pairs (**b**) and RNA unique molecular identifiers (UMIs; **c**). Cells are classified as human, mouse or doublets on the basis of species-specific reads. **d,** Comparison of chromosome 3 contact maps and A/B compartment profiles between HSPC ChromE-seq and published bulk HSPC Hi-C^22^. **e,** Single-cell HSPC ChromE-seq contact profiles across the *CD34* locus for 500 representative cells, with the corresponding aggregated contact map shown above. **f,g,** Comparison of distal Hi-C contact (intrachromosomal contact pairs separated by more than 1 kb) recovery (**f**) and RNA detection (**g**) between ChromE-seq and published joint chromatin-conformation/transcriptome assays^12–16^, and ChromE-seq datasets generated from HEK293 cells, HSPCs and brain. Box plots show the median and interquartile range (25th-75th percentiles), with whiskers extending to 1.5× the interquartile range. **h,** Estimated cellular throughput per assay across methods. **i,** Estimated library preparation cost per 1,000 cells across methods.

**Extended Data Figure 2.**
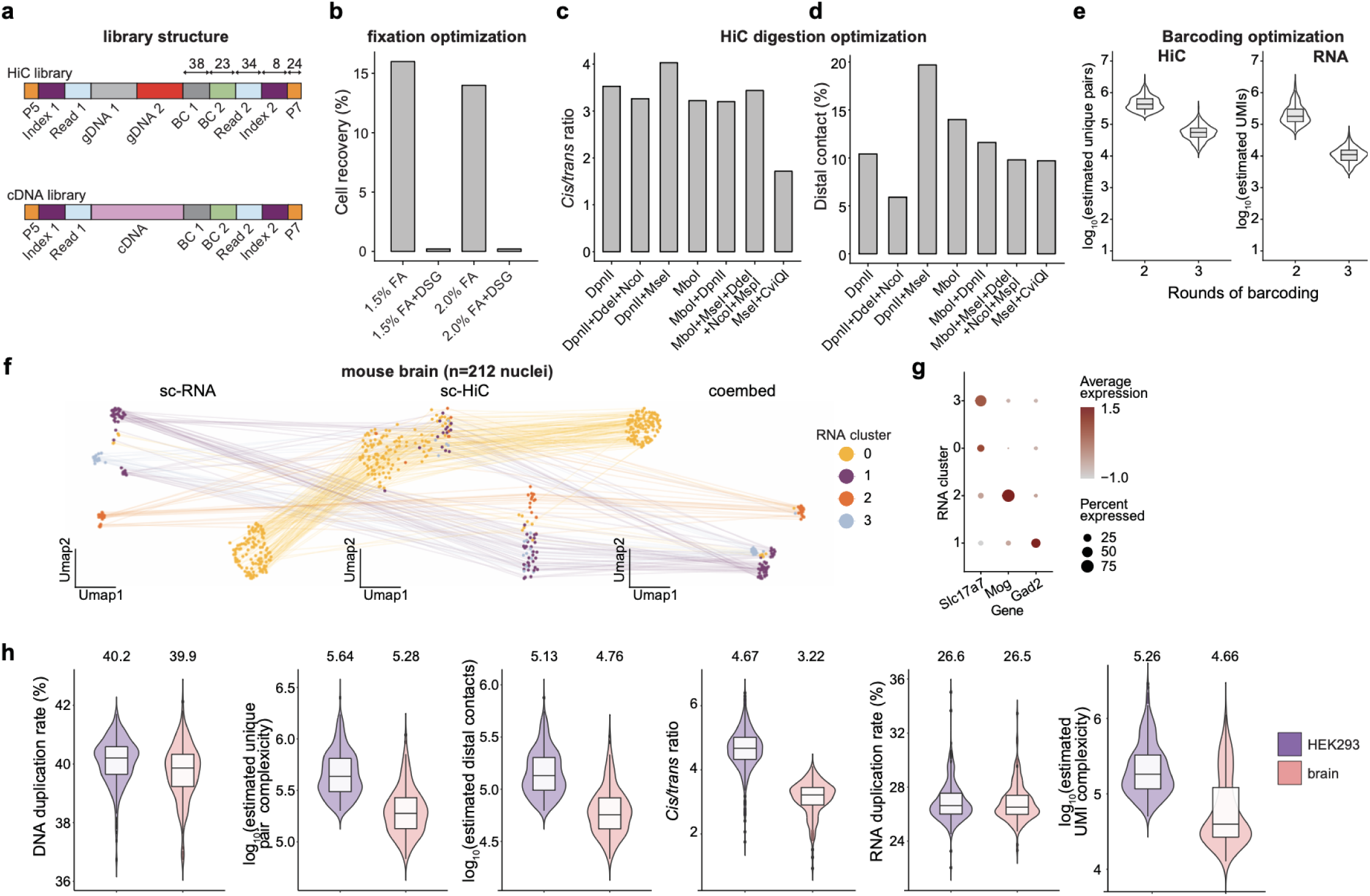
Optimization and validation of ChromE-seq. **a,** Schematic of Hi-C and cDNA library structures generated by ChromE-seq, including barcode and index positions. **b,** Optimization of fixation conditions, showing cell recovery following formaldehyde fixation with or without DSG. **c,d,** Optimization of restriction-enzyme conditions for Hi-C, assessed by *cis/trans* contact ratio (**c**) and proportion of distal chromatin contacts (**d**). **e,** Comparison of two versus three rounds of combinatorial barcoding, showing estimated Hi-C unique pairs and RNA UMIs per cell. Box plots show the median and interquartile range (25th-75th percentiles), with whiskers extending to 1.5× the interquartile range. **f,** Joint RNA and Hi-C profiling of mouse brain nuclei. UMAP embeddings are shown for RNA, Hi-C and the combined representation, with lines linking measurements from the same cell. **g,** Expression of representative neuronal marker genes across RNA-defined clusters. Dot size indicates the fraction of cells expressing each gene and colour indicates average expression. **h,** ChromE-seq performance metrics in HEK293 cells and mouse brain nuclei, including DNA duplication rate, unique-pair complexity, distal contacts, *cis/trans* ratio, RNA duplication rate, and estimated UMI complexity. Numbers above the violins indicate median values. Box plots show the median and interquartile range (25th-75th percentiles), with whiskers extending to 1.5× the interquartile range.

**Extended Data Figure 3.**
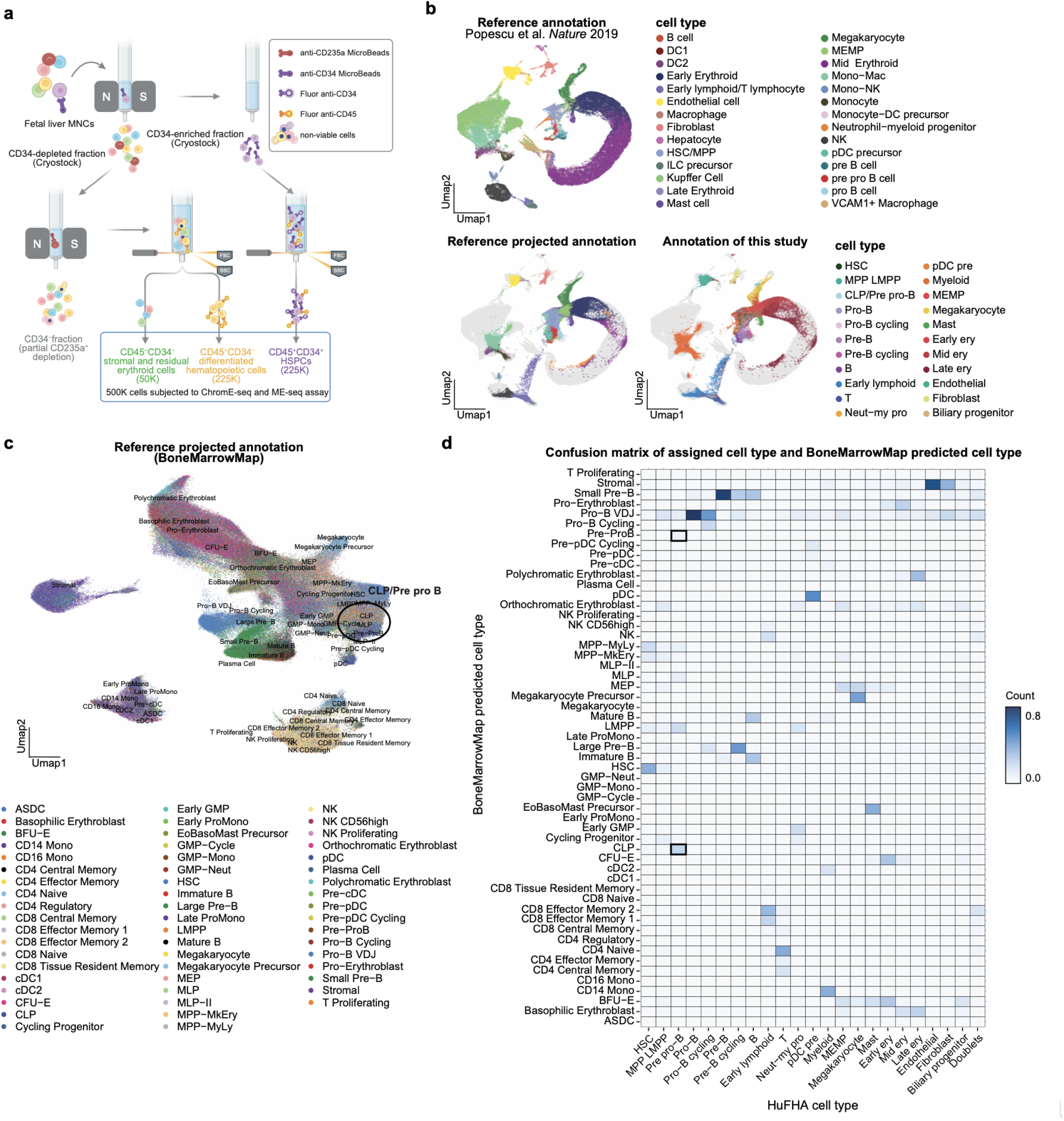
Sample processing and cell-type annotation of the fetal liver atlas. **a,** Experimental workflow for enriching CD34^+^ HSPCs, CD45^+^ immune cells and CD45^−^ stromal cells from dissociated fetal liver samples. Fetal liver mononuclear cells (MNCs) were separated by magnetic CD34 enrichment and cryopreserved as CD34-enriched and CD34-depleted fractions. The CD34-depleted fraction was subsequently depleted of CD235a^+^ mature erythroid cells with magnetic beads and sorted into CD45^−^CD34^−^ stromal and residual erythroid cells and CD45^+^CD34^−^ differentiated immune cells, while CD45^+^CD34^+^ HSPCs were sorted from the CD34-enriched fraction. **b,** Cell-type annotation of the fetal liver atlas using reference mapping to a human fetal liver dataset (Popescu et al. *Nature*, 2019)^23^. Reference annotations, projected labels and final annotations used in this study are shown. **c,** Independent projection of atlas cells onto BoneMarrowMap (Zeng et al. *Blood Cancer Discovery*, 2025)^24^, showing predicted haematopoietic cell identities across the integrated embedding. **d,** Confusion matrix comparing HuFHA cell-type annotations with BoneMarrowMap-predicted identities. Colour indicates the fraction of cells assigned to each reference cell type.

**Extended Data Figure 4.**
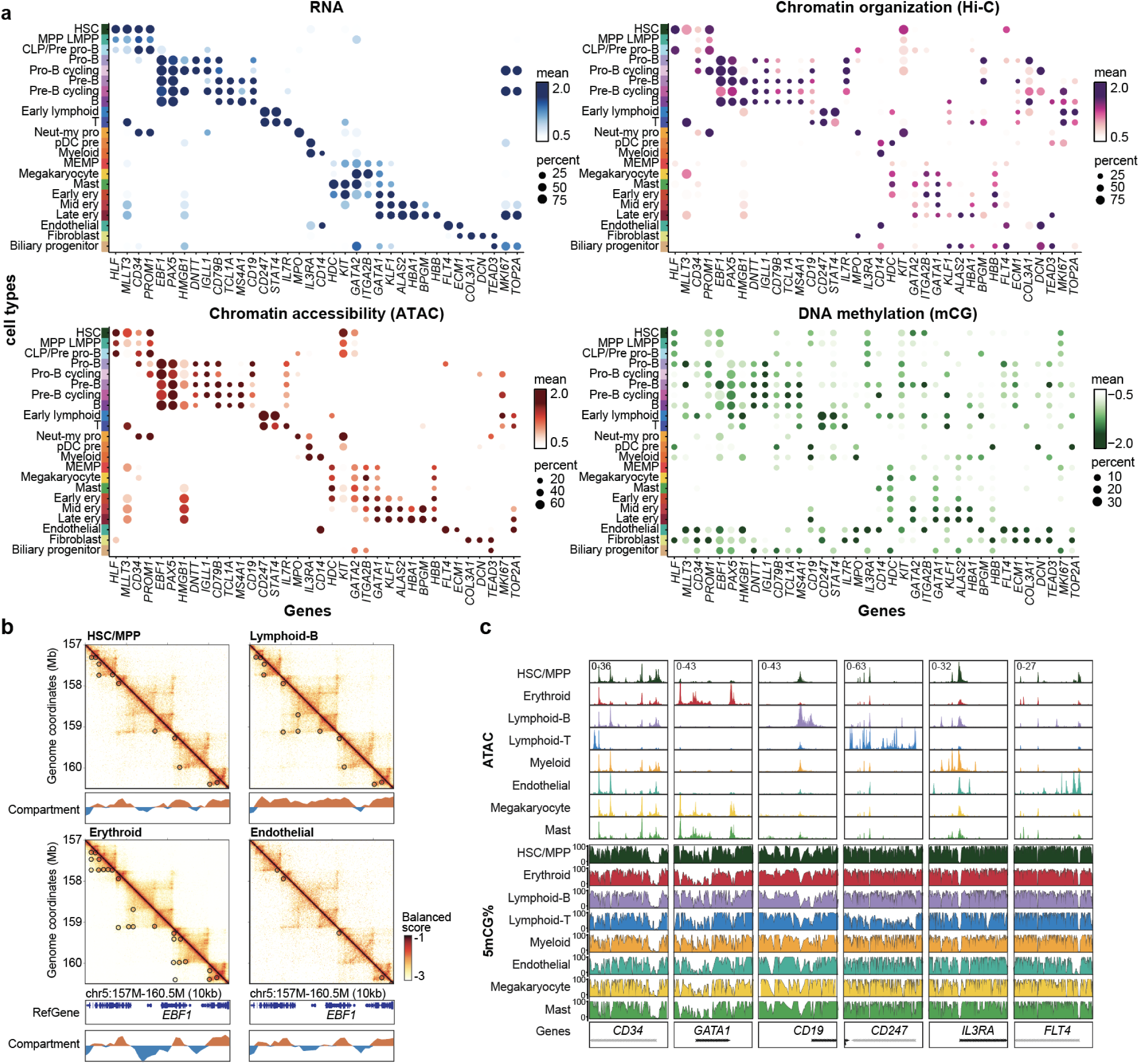
Multimodal features of fetal liver cell identity. **a,** Expression and epigenetic profiles of canonical lineage-marker genes across annotated cell types. Dot plots show RNA expression, Hi-C contact signal, chromatin accessibility, and DNA methylation; dot size indicates the fraction of cells with detectable signal and colour indicates mean normalized signal. **b,** Pseudobulk Hi-C contact maps across the *EBF1* locus in HSC/MPP, B-lymphoid, erythroid and endothelial populations. A/B compartment profiles and cell-type-specific chromatin loops are shown. **c,** Pseudobulk chromatin-accessibility and DNA-methylation profiles across representative lineage-marker loci in major haematopoietic lineages.

**Extended Data Fig. 5.**
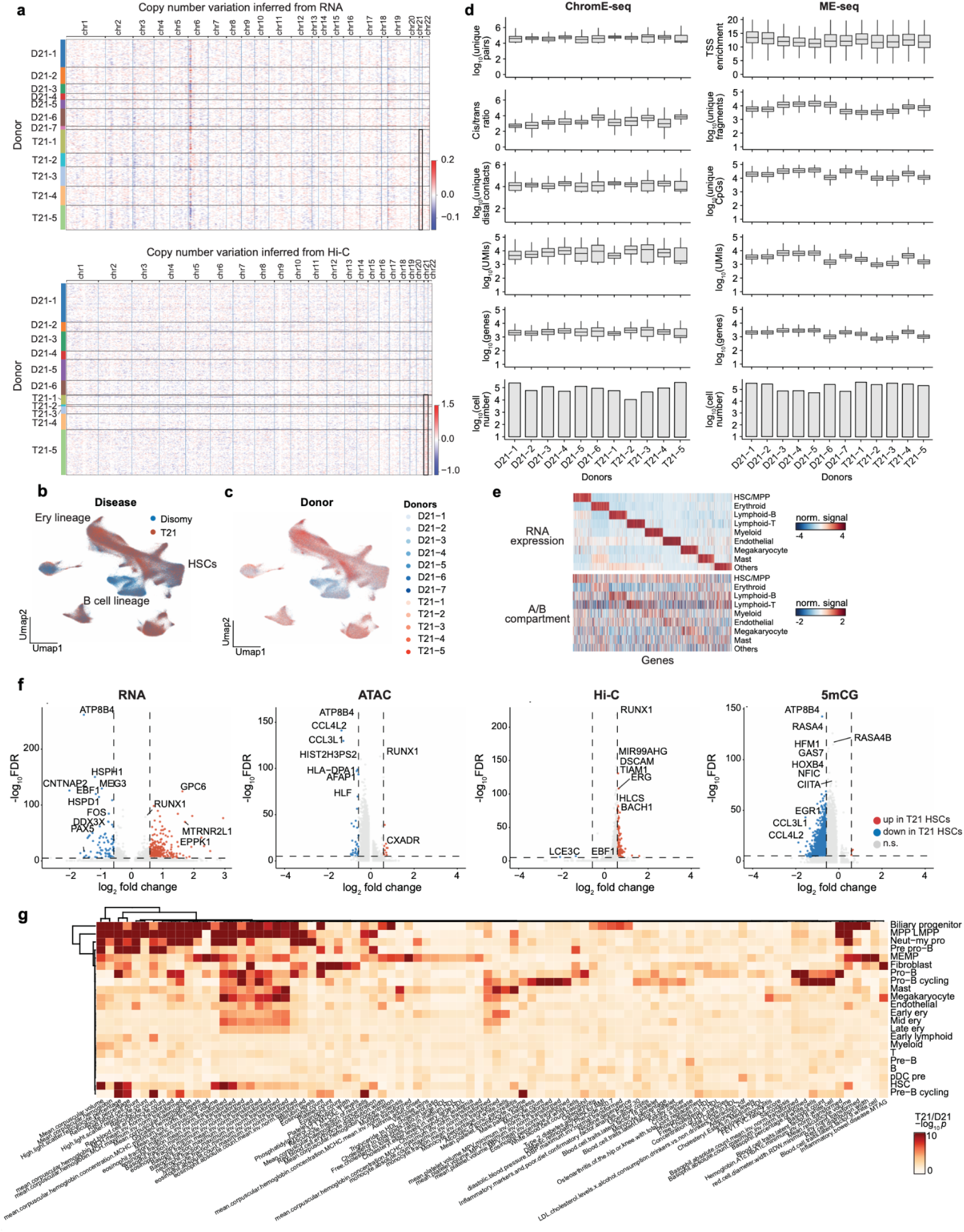
Multimodal characterization of disomic and trisomy 21 fetal haematopoiesis. **a,** Copy-number profiles inferred independently from RNA (top) and Hi-C (bottom) across donors. Chromosome 21 is highlighted, showing the expected copy-number increase in T21 samples. **b,c,** Integrated single-cell UMAPs coloured by trisomy status (**b**) and donor identity (**c**), showing representation of disomic and T21 donors across major haematopoietic populations. **d,** Donor-level quality-control metrics for ChromE-seq and ME-seq, including Hi-C unique pairs, cis/trans ratio and distal contacts, RNA UMIs and detected genes, transcription-start-site enrichment, ATAC fragments, CpG coverage and total cell recovery. **e,** Normalized RNA expression (top) and A/B compartment signal (bottom) across variable lineage-associated genes in major lineages. **f,** Differential RNA expression, chromatin accessibility, Hi-C contacts, and DNA methylation between T21 and disomic HSC/MPP. **g,** Cell-type-specific enrichment of T21-associated genetic variants. Colour indicates enrichment significance.

**Extended Data Figure 6.**
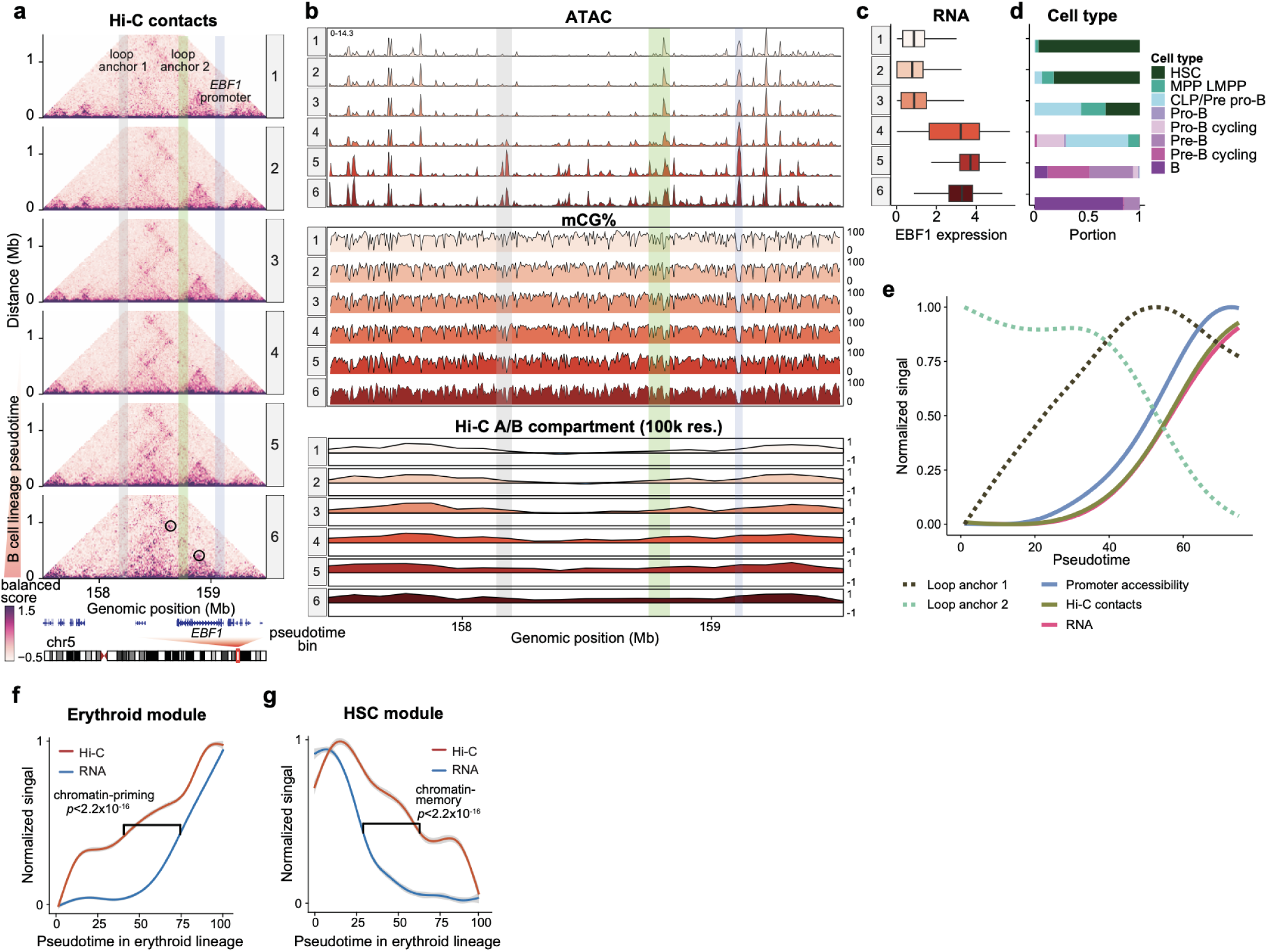
Staged regulatory remodeling of the *EBF1* locus during B-lineage differentiation. **a,** Pseudobulk Hi-C contact maps across the *EBF1* locus in six B-lineage pseudotime bins. The *EBF1* promoter and two distal loop anchors are highlighted. **b,** Chromatin accessibility, CpG methylation, and A/B compartment profiles across the *EBF1* locus for the same pseudotime bins. **c,** *EBF1* RNA expression across B-lineage pseudotime bins. **d,** Cell-type composition of each pseudotime bin, showing progression from HSC/MPP through CLP/pre-pro-B and downstream B-lineage states. **e,** Smoothed normalized signals for the two distal loop anchors, *EBF1* promoter accessibility, overall Hi-C contact signal and *EBF1* RNA expression along B-lineage pseudotime. **f,g,** Normalized Hi-C and RNA signals along erythroid pseudotime for an erythroid-associated module (**f**) and an HSC-associated module (**g**), illustrating chromatin priming and persistence, respectively. *K-S* test.

**Extended Data Figure 7.**
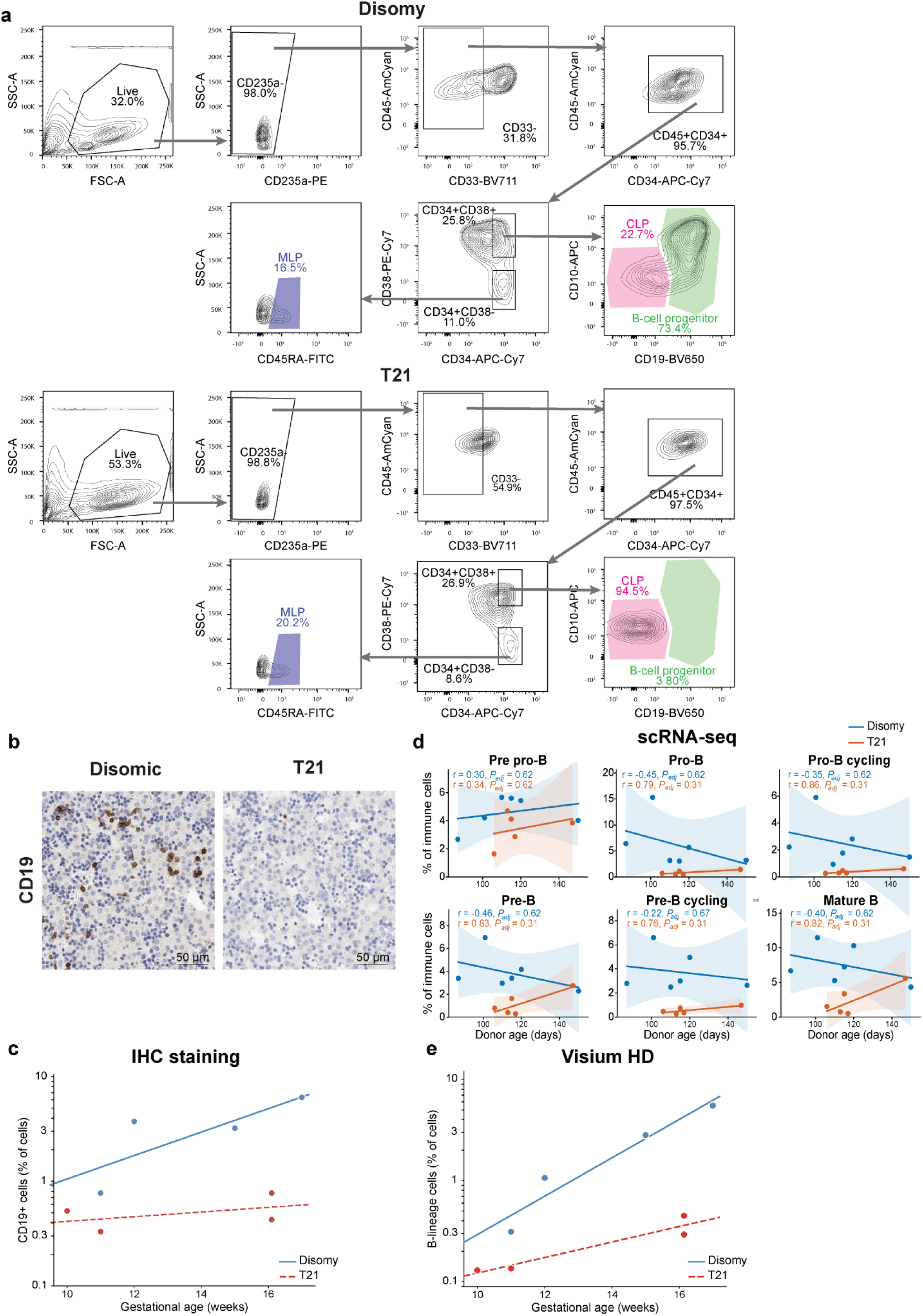
B-cell progenitors are depleted in trisomy 21 fetal liver. **a,** Representative flow-cytometry gating strategy for disomic and T21 fetal liver samples, identifying HSC/MPP, MLP, CLP, and downstream B-cell progenitors **b,** Representative CD19 immunohistochemistry in disomic and T21 fetal liver sections. Scale bars, 50 μm. **c,** Quantification of the percentage of CD19⁺ cells in IHC-stained disomic and T21 fetal liver sections as a function of gestational age. Points represent individual specimens, and lines indicate linear regression. **d,** Abundance of B-lineage populations inferred from scRNA-seq as a function of donor gestational age in disomic and T21 fetal liver. Points represent individual donors; lines indicate linear regression fits and shaded regions indicate 95% confidence intervals. Pearson correlation coefficients (*r*) and associated *P* values are shown. **e,** B-lineage cell abundance estimated from Visium HD spatial transcriptomics in the same fetal liver specimens. Points represent individual specimens and lines indicate linear regression fits.

**Extended Data Figure 8.**
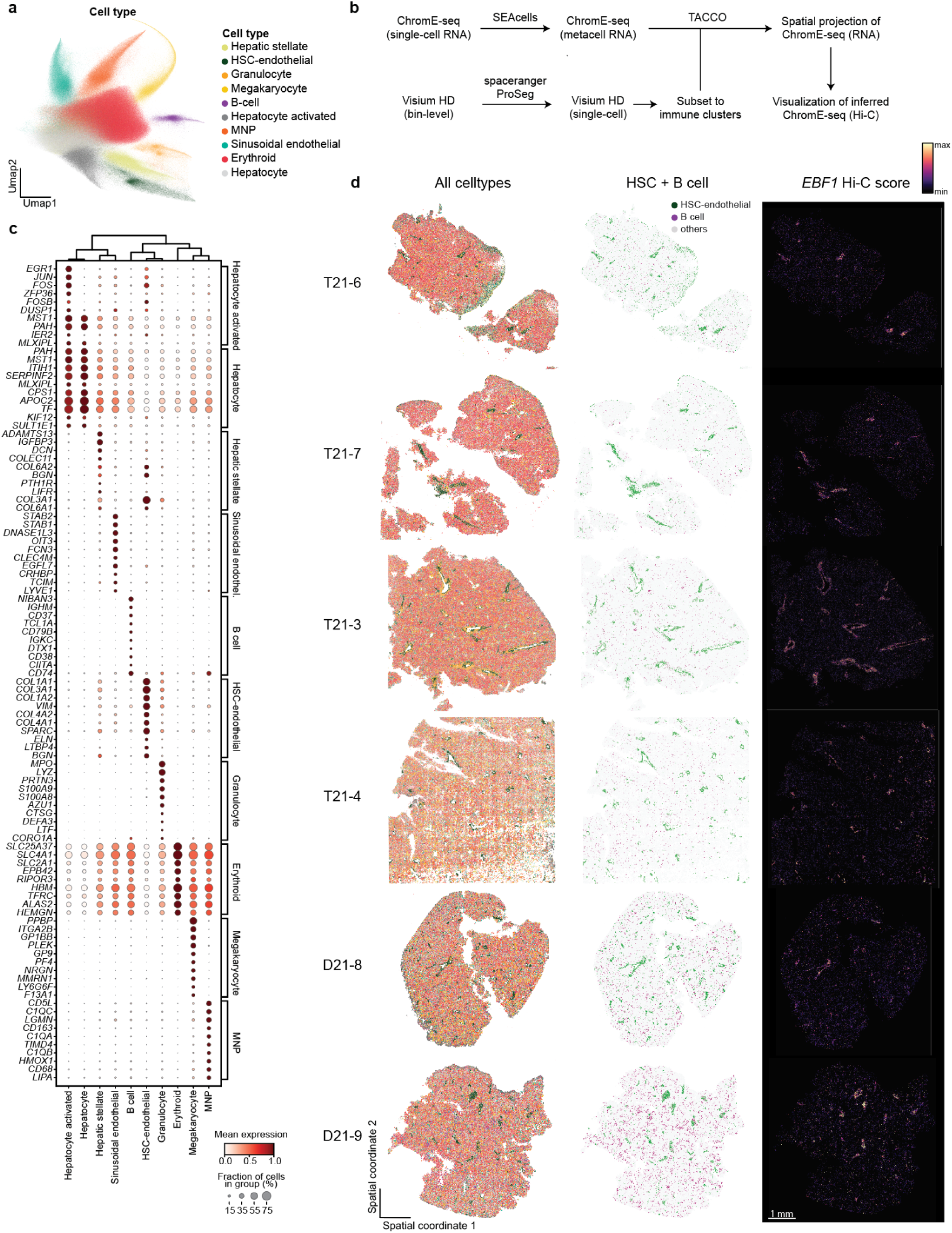
Spatial projection of inferred *EBF1* chromatin-contact signal in fetal liver. **a,** UMAP embedding of Visium HD profiles from fetal liver samples, coloured by annotated cell type. **b,** Schematic of the integration workflow used to project ChromE-seq-derived information onto Visium HD data. ChromE-seq single-cell RNA profiles were first aggregated into metacells with SEACells, and Visium HD bin-level profiles were processed into single-cell profiles with spaceranger ProSeg. Cell-state annotations were transferred by TACCO, enabling spatial projection of ChromE-seq RNA states and visualization of inferred ChromE-seq Hi-C features. **c,** Dot plot showing representative marker genes for annotated Visium HD cell types. Dot size indicates the fraction of cells in each group expressing the gene, and colour indicates mean expression. **d,** Spatial maps for each Visium HD sample showing all annotated cell types (left), HSC-endothelial and B-cell populations highlighted against other cells (middle), and the projected *EBF1* Hi-C gene score (right).

**Extended Data Figure 9.**
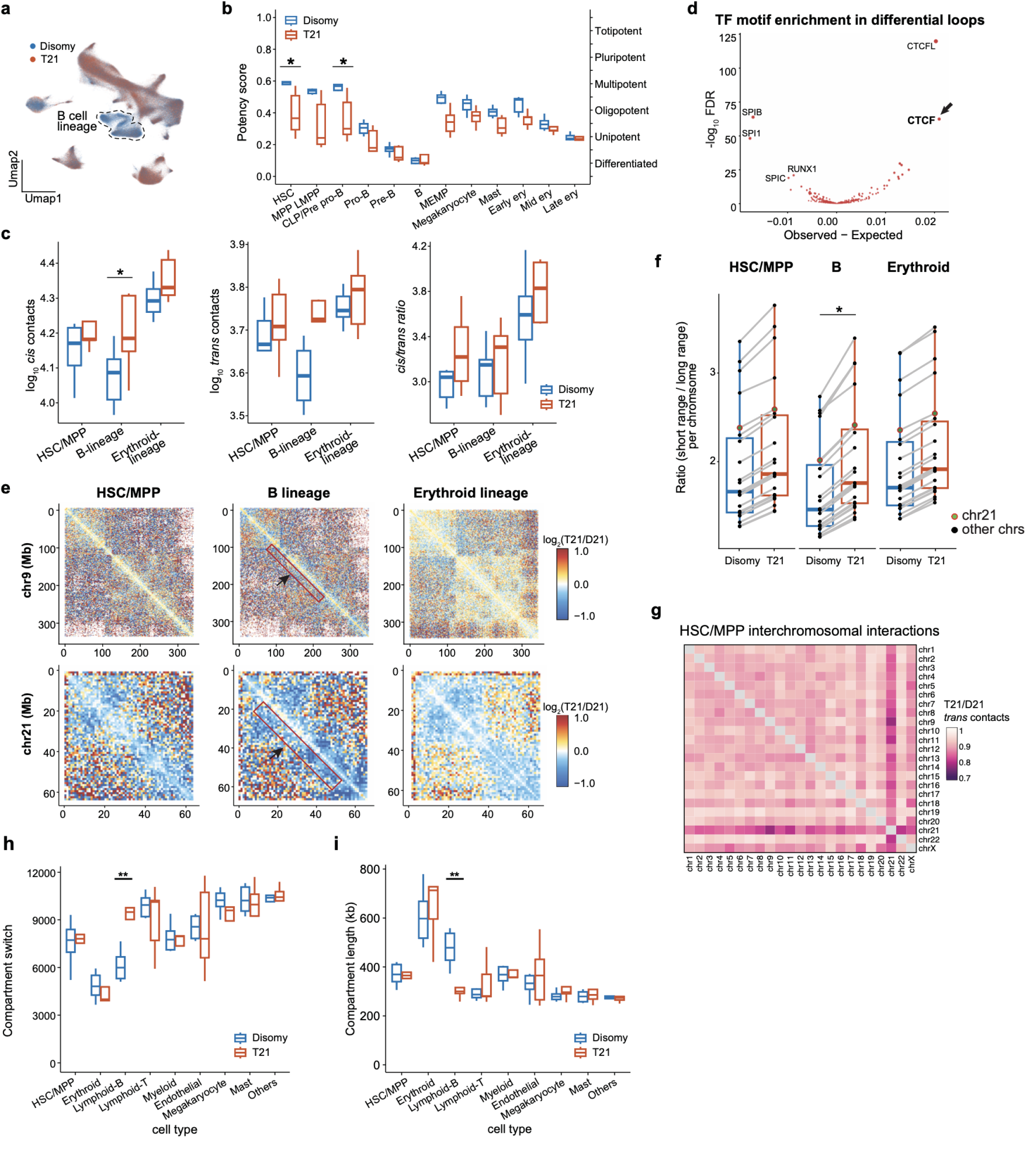
Trisomy 21 alters developmental potency and higher-order chromatin organization. **a,** Single-cell UMAP coloured by trisomy status, with the B-lineage compartment indicated. **b,** CytoTRACE2 developmental-potency scores projected onto the UMAP embedding. **c,** Global chromatin-contact properties across HSC/MPP, B-lineage and erythroid cells in disomy and T21, including *cis* contacts, *trans* contacts and the *cis/trans* contact ratio. **d,** Transcription factor (TF) motif enrichment within T21-differential chromatin loops. Top enriched regulators are indicated. **e,** T21-to-disomy Hi-C contact-ratio maps for chromosome 9 (top) and chromosome 21 (bottom) in HSC/MPP, B-lineage and erythroid cells. Red indicates increased and blue decreased contact frequency in T21 relative to disomy. **f,** Ratio of short- to long-range chromatin interactions in HSC/MPP, B-lineage and erythroid cells, shown for individual chromosomes in disomic and T21 samples. Chromosome 21 is highlighted. **g,** Chromosome-pair matrix showing T21-to-disomy ratios of interchromosomal contacts in HSC/MPP. **h,** Number of A/B compartment switches across major cell types in disomic and T21 samples. **i,** A/B compartment length across major cell types in disomic and T21 samples. * *p* < 0.05, ** *p* < 0.01, *t*-test.

**Extended Data Figure 10.**
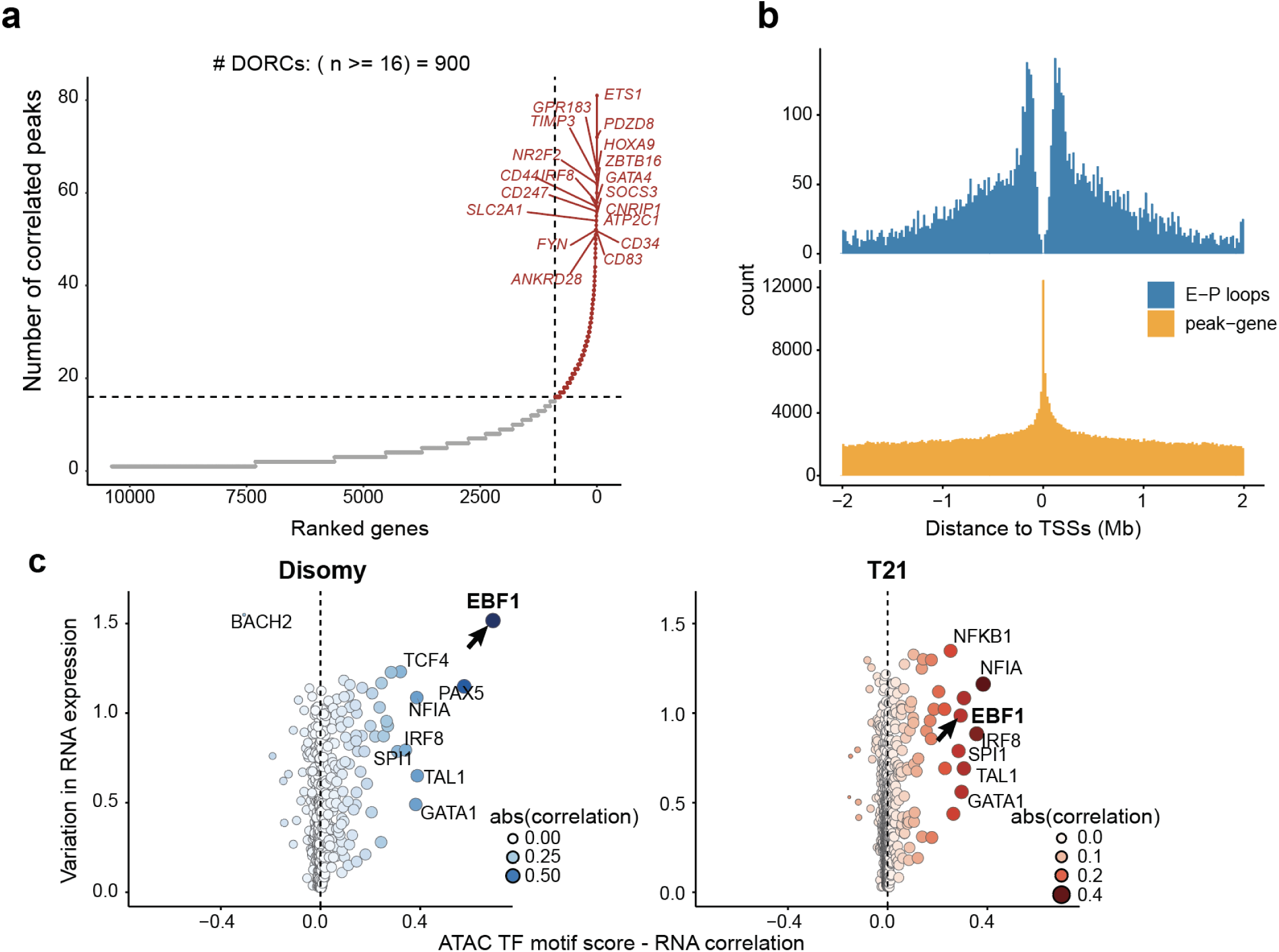
Multimodal regulatory coupling identifies EBF1 as a highly connected B-lineage regulator. **a,** Ranked distribution of DORC genes by number of correlated regulatory peaks. Dashed lines indicate the threshold used to define DORCs (≥16 correlated peaks), identifying 900 DORC genes. **b,** Genomic-distance distributions of enhancer–promoter Hi-C loops and peak–gene associations relative to transcription start sites. **c,** Relationships between ATAC TF-motif score and RNA expression in disomic and T21 cells. Point size and colour indicate the absolute correlation strength.

**Extended Data Figure 11.**
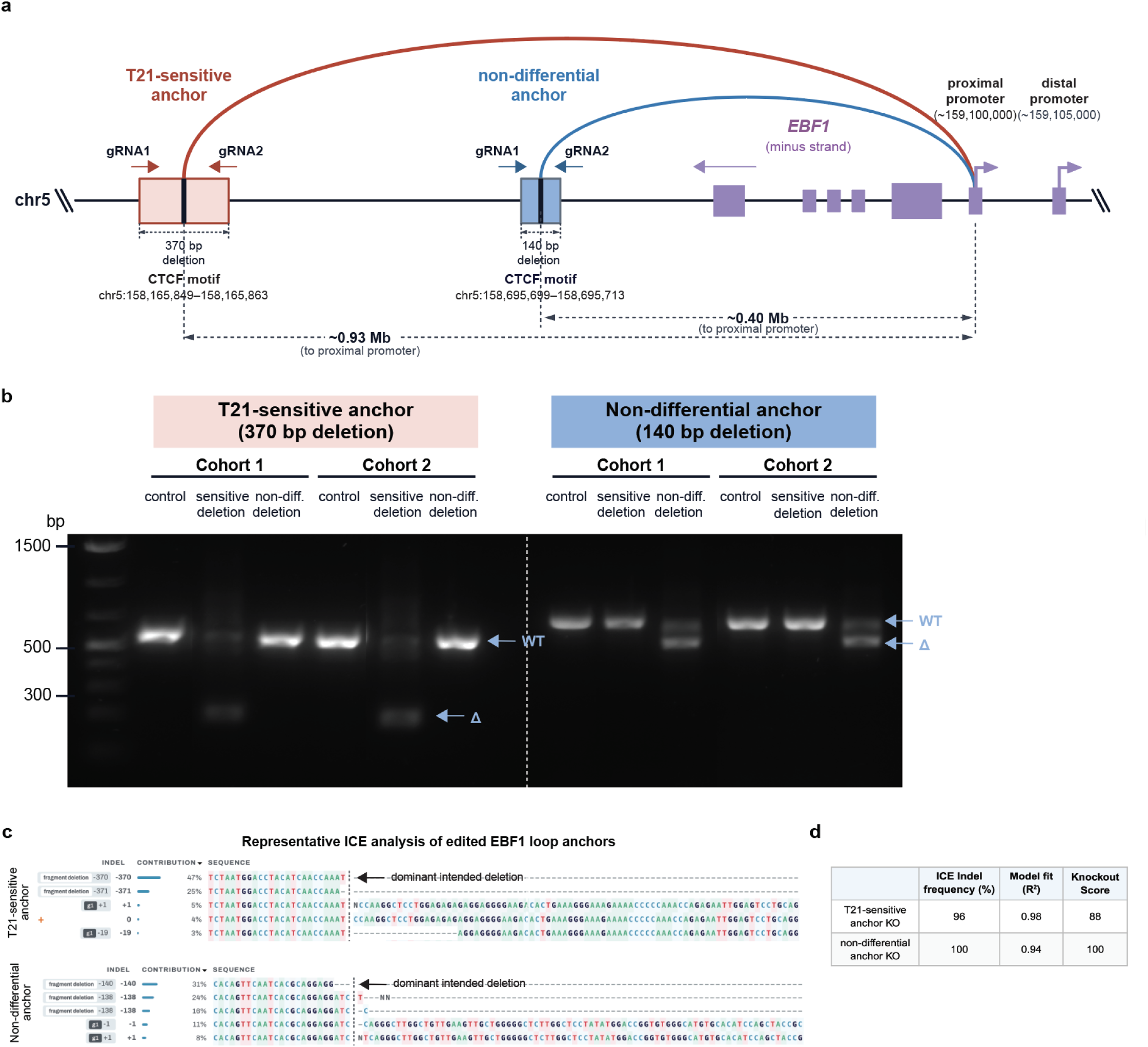
Validation of CRISPR editing at *EBF1* loop anchors. **a,** Schematic of the CRISPR/Cas9 strategy used to disrupt the T21-sensitive and non-differential *EBF1* loop anchors. The locations of the two anchors relative to the *EBF1* gene body and proximal and distal promoters are shown. **b,** PCR genotyping of edited human fetal liver HSPCs from two independent cohorts immediately before xenotransplantation. The T21-sensitive and non-differential anchor regions were amplified separately to detect wild-type (WT) and deletion (Δ) products. Expected WT and deletion products are indicated by arrows. **c,** Representative ICE analysis of the edited *EBF1* loop anchors following Sanger sequencing. The top five most abundant inferred sequence outcomes are shown. **d,** Summary of ICE-estimated editing metrics for representative T21-sensitive and non-differential anchor knockout samples.

## Extended data tables

**Extended Data Table 1. ChromE-seq optimization conditions.**

**Extended Data Table 2. Metadata for human–mouse species-mixing experiments.**

**Extended Data Table 3. Comparison of ChromE-seq with published joint chromatin conformation and transcriptome methods.**

**Extended Data Table 4. ChromE-seq cost breakdown.**

**Extended Data Table 5. Fetal liver donor information.**

**Extended Data Table 6. Differential Hi-C loops between T21 and disomy across cell types.**

**Extended Data Table 7. ChromE-seq oligo design.**

**Extended Data Table 8. ChromE-seq index primers.**

**Extended Data Table 9. Antibodies used for flow-cytometric analysis of human fetal liver and xenograft samples.**

**Extended Data Table 10. gRNA sequences.**

## Methods

### Animal studies

All mouse experiments were approved by the Icahn School of Medicine at Mount Sinai Institutional Animal Care and Use Committee. We confirm that all experiments conform to the relevant regulatory and ethical standards. All xenotransplantations were performed in 8 to 12-week-old *NOD.Cg-Prkdc^scid^Il2rg^tm1Wjl^/SzJ* (NSG) mice (JAX) that were sublethally irradiated with 230 cGy, 24 hours before transplantation.

### Human samples

Human fetal liver samples were obtained from elective pregnancy terminations from the Developmental Origins of Health and Disease (DOHaD) Biorepository at the Icahn School of Medicine at Mount Sinai with written informed consent following guidelines approved by the Icahn School of Medicine at Mount Sinai Institutional Review Board. Fetal liver samples were collected between 10 to 21 weeks’ gestation.

### Cell culture

HEK293 cells (ATCC, CRL-1573) were grown in DMEM (high glucose; Gibco) with 10% heat-inactivated FBS and 1% penicillin–streptomycin, at 37 °C in 5% CO_2_. Cells were split at 70-90% confluence using TrypLE Express (12604021, ThermoFisher) and used between passages 4 and 10. Cells were digested with TrypLE Express and resuspended in PBSI (containing 0.25% RNase inhibitor, Y9240L, QIAGEN; and 0.04% bovine serum albumin, BSA, A8412, Millipore Sigma) for preparing single-cell suspensions.

### Mouse brain sample processing

C57BL/6J mice were euthanized by CO_2_ inhalation followed by cervical dislocation. Whole brains were removed quickly, snap-frozen in liquid nitrogen, and stored at −80 °C. For experiments, the prefrontal cortex (PFC) was dissected from the frozen brains on a chilled plate and processed as previously described^17^ for ChromE-seq. Frozen tissue was placed into a pre-chilled 2 mL Dounce homogenizer containing 430 μL of 1× homogenization buffer (320 mM sucrose, 0.1 mM EDTA, 0.1% NP-40, 5 mM CaCl_2_, 3 mM Mg(Ac)_2_, 10 mM Tris-HCl pH 8.0, 17 μM PMSF, 0.17 mM DTT, and 0.25% RNase inhibitor, Y9240L, QIAGEN). The tissue was homogenized with 10 strokes using pestle A, followed by 20 strokes with pestle B. Next, 400 μL of the homogenate was transferred to a pre-chilled 2 mL graduated microcentrifuge tube. An equal volume (400 μL) of 50% iodixanol solution (prepared in 1× homogenization buffer) was added and mixed to create a 25% iodixanol layer. Subsequently, 600 μL of 29% iodixanol solution (160 mM sucrose in 1× homogenization buffer) was carefully layered beneath the 25% iodixanol layer, followed by 600 μL of 35% iodixanol solution (160 mM sucrose in 1× homogenization buffer) layered at the bottom. Nuclei were isolated by centrifugation at 3,000 g for 20 minutes at 4°C using a swinging-bucket rotor. The resulting nuclear pellet was resuspended in HCBI buffer (0.04% BSA, 10 mM NaCl, 3 mM MgCl_2_, and 10 mM HEPES, pH 7.2, 0.25% RNase inhibitor, Y9240L, QIAGEN) and immediately processed for fixation.

### Fetal liver processing

Fresh fetal liver tissue was finely minced with razor blades and divided between two 50 mL tubes, each containing 40 mL prewarmed IMDM, 5 mL collagenase IV stock solution (1 mg/mL; STEMCELL Technologies), and 25 μL DNase I (10 mg/mL; Roche). Samples were incubated at 37 °C for 30 min with rotation. Dissociated tissue was filtered through a 40 μm cell strainer (Thermo Fisher Scientific), and residual tissue was gently passed through the strainer using the rubber plunger of a 5 mL syringe (BD Biosciences). Cells were centrifuged at 350 × *g* for 10 min.

Erythrocytes were lysed with 5 mL ammonium chloride solution (STEMCELL Technologies) per tube for 5 min at room temperature. Lysis was quenched with IMDM, and cells were centrifuged at 350 × *g* for 10 min. Cell pellets were pooled and resuspended in autoMACS Running Buffer (Miltenyi Biotec). CD34^+^ cells were enriched using the human CD34 MicroBead Kit (Miltenyi Biotec) according to the manufacturer’s instructions, using approximately one LS column (Miltenyi Biotec) per gram of starting tissue. The CD34-enriched positive fraction and CD34-depleted flow-through were collected separately and cryopreserved in 90% FBS (Cytiva) and 10% dimethyl sulfoxide (DMSO; Sigma-Aldrich) in liquid nitrogen until further use.

### Isolation of fetal liver cell population by fluorescence activated cell sorting

For isolation of CD34^+^ HSPCs, cryopreserved CD34-enriched fractions were thawed by slow, dropwise addition of thawing medium consisting of X-VIVO 10 (Lonza), 50% FBS (Cytiva) and DNase I (100 μg/mL; Roche). Cells were centrifuged at 350 × *g* for 10 min and resuspended in PBS containing 2.5% FBS. Cryopreserved CD34-depleted fractions were processed in parallel for isolation of CD45^+^ and CD45^−^ populations. Because the CD34-depleted fraction contained a high abundance of erythroid cells, CD235a^+^ cells were partially depleted by magnetic separation using anti-CD235a magnetic microbeads (Miltenyi Biotec) before fluorescence-activated cell sorting. This depletion was performed to reduce erythroid overrepresentation rather than to obtain a completely CD235a-negative population, and residual CD235a^+^ cells remained in the resulting fraction.

Cells were resuspended at approximately 5 × 10^6^ cells/mL and stained for 60 min at 4 °C with CD34, CD45, and CD235a PE. Antibody clones, fluorochromes, dilutions and suppliers are provided in **Extended Data Table 9**. Viability Stain 450 (1:1,000; BD Biosciences) was added immediately before sorting. Viable singlets were sorted into three populations: CD34^+^ HSPCs from the CD34-enriched fraction, and CD34^−^CD45^+^ differentiated immune cells and CD34^−^CD45^−^ stromal cells from the CD34-depleted fraction. CD235a staining was used to characterize the erythroid composition of the CD45^−^ fraction but was not used as a sorting criterion. Cells were collected into PBS (containing 10% FBS and 0.25% RNase inhibitor, Y9240L, QIAGEN).

For ChromE-seq, target sorting yields were approximately 5 × 10^5^ CD34^+^ cells, 5 × 10^5^ CD45^+^ cells and 1 × 10^5^ CD45^−^ cells per sample. For ME-seq, target yields were approximately 3.5 × 10^5^ CD34^+^ cells, 3.5 × 10^5^ CD45^+^ cells and 5 × 10^4^ CD45^−^ cells per sample. Where cell availability permitted, additional cells were collected. Cells were sorted on a BD FACSMelody cell sorter (BD Biosciences). Immediately after sorting, cells were fixed according to the corresponding downstream ChromE-seq or ME-seq protocol.

### ChromE-seq sample processing

For the human–mouse mixing experiment, HEK293 cells in suspension were centrifuged at 300 × g for 5 min and resuspended in PBSI at 1 × 10^6^ cells/mL. Cells were fixed with 1.5% formaldehyde (28906, Thermo Fisher Scientific) for 5 min at room temperature. Fixation was quenched by adding 120 μL of Quench buffer (1.17 M glycine, 0.42 M Tris-HCl, pH 8.0, and 0.86% BSA) per milliliter of the original cell suspension. Samples were incubated on ice for 10 min and centrifuged at 750 × g for 5 min. The supernatant was removed, and the cell pellet was washed twice with 500 μL of PBSI. To isolate nuclei, cells were resuspended in 100 μL of NI-IGEPAL buffer (10 mM Tris pH 7.5, 10 mM NaCl, 3 mM MgCl_2_, 0.2% IGEPAL-630, freshly added 0.1 U/μL Enzymatics RNase Inhibitor, and 0.05 U/μL SUPERase RI), and incubated on ice for 5 min. Immediately after incubation, 900 μL of NI buffer was added to limit further permeabilization. Mouse brain nuclei suspended in HCBI buffer were fixed with 1.0% formaldehyde for 5 min, quenched by adding 120 μL of the same Quench buffer per milliliter of the original nuclei suspension, and incubated on ice for 10 min. Mouse brain nuclei were washed twice with 500 μL of HCBI buffer and pooled with permeabilized HEK293 nuclei at a 1:3 mouse-to-human ratio. The pooled nuclei were then processed according to the ChromE-seq protocol, beginning at the Nucleosome Depletion step. For fetal liver samples, the three sorted cell populations (CD34^+^ cells, CD45^+^ cells, and CD45^−^ cells) were pooled and processed directly according to the ChromE-seq protocol.

### ChromE-seq protocol

ChromE-seq combines *in situ* proximity ligation for capturing chromatin contacts with parallel reverse transcription for mRNA capture, followed by combinatorial indexing to assign both modalities to individual cells. Briefly, cells or nuclei were fixed to preserve native chromatin interactions and RNA molecules, followed by permeabilization, nucleosome depletion, and *in situ* chromatin digestion using an optimized restriction enzyme combination (DpnII and MseI). Digested chromatin ends were proximity-ligated to generate Hi-C contacts. In parallel, mRNA was reverse-transcribed using biotinylated poly-dT and random primers to preserve transcriptome information from the same cells. Nuclei were then subjected to split-pool combinatorial indexing, in which chromatin contacts and cDNA molecules were sequentially barcoded across two rounds to generate cell-specific barcode combinations. After reverse crosslinking, chromatin and cDNA fractions were separated, amplified, tagmented using Tn5, and converted into sequencing libraries for paired analysis of single-cell chromatin conformation and RNA expression. The ChromE-seq libraries were sequenced on a NovaSeq using standard primers with paired-end 150 bp reads and 8 bp dual indexes (Read 1: 150 cycles; Index 1: 8 cycles; Index 2: 8 cycles; Read 2: 150 cycles). Sequences of the split-pool barcoding oligonucleotides and PCR index primers are provided in **Extended Data Tables 7 and 8**, respectively. A detailed step-by-step protocol is available at Protocols.io (dx.doi.org/10.17504/protocols.io.e6nvwbq4wvmk/v1).

### ME-seq sample processing

Following isolation from fetal liver, the three sorted cell populations (CD34⁺, CD34⁻CD45⁺, and CD34⁻CD45⁻ cells) were centrifuged at 300 × g for 5 min and resuspended in PBSI at 1 × 10^6^ cells/mL. Cells were fixed with 1.0% formaldehyde for 5 min at room temperature. Fixation was quenched by adding 120 μL of Quench buffer per milliliter of the original cell suspension, followed by incubation on ice for 10 min. Samples were centrifuged at 750 × g for 5 min, the supernatant was removed, and the cell pellet was washed twice with 500 μL of PBSI. Nuclei were isolated by resuspending cells in 100 μL of NI-NP-40 buffer (10 mM Tris buffer pH 7.5, 10 mM NaCl, 3 mM MgCl_2_, 0.1% NP-40, freshly added 0.1 U/μL Enzymatics RNase Inhibitor, and 0.05 U/μL SUPERase RI), and incubated on ice for 3 min. Immediately after incubation, 900 μL of NI buffer was added to limit further permeabilization. Samples were subsequently processed according to the step-by-step ME-seq protocol on Protocols.io (dx.doi.org/10.17504/protocols.io.5qpvowmwxl4o/v1), beginning at the Transposition step.

### Visium HD sample processing

A small piece of fetal liver tissue was fixed in 10% neutral-buffered formalin (Thermo Fisher Scientific) for 24 hours and subsequently processed and embedded in paraffin. Formalin-fixed, paraffin-embedded (FFPE) tissue blocks were sectioned onto glass microscope slides with 5 µm thickness. Sequencing-based spatial transcriptomics was then performed using the 10x Genomics Visium HD Spatial Gene Expression platform following manufacturer guidelines for FFPE samples (10x User Guide CG000685). High-resolution H&E images were captured after deparaffinization and decrosslinking, prior to transcript probe hybridization. Visium CytAssist Spatial Gene Expression Slides containing 6.5 mm x 6.5 capture areas were used. After spatial transcriptomics processing and library construction, spatial transcriptomics samples were sequenced on an Illumina NovaSeq X through Azenta.

### ChromE-seq quality control, pre-processing, and visualization

DNA sequencing reads were aligned to hg38 or mm10 reference genome using BWA-MEM2 (v2.2.1) with argument -SP5M specified, and alignments were converted to BAM format using SAMtools (v1.11). Chromatin contacts were extracted from aligned reads using pairtools (v1.0.2), with hg38 chromosome sizes provided for coordinate parsing and mapping quality scores retained. To preserve multi-ligation information, contacts were parsed using the --walks-policy all option with a minimum mapping quality of 1, while retaining the original pair orientation.

Multi-ligated reads were processed using a custom walk-rescue procedure, based on a previously published single-cell Hi-C workflow^12^. Read identifiers were simplified to retain molecule-associated indexing information, and records sharing the same read identifier were grouped using a rescue_walk.py script^12^. For each molecule, uniquely mapped genomic ends were identified and all possible pairwise combinations of these ends were evaluated. Contacts that had already been directly observed were retained, while additional pairwise contacts between mapped ends that were not directly observed were generated to recover contacts from multi-ligation events. Unmapped ends and self-pairs were excluded from contact reconstruction. The rescued contacts were subsequently canonicalized according to genomic coordinates and sorted using pairtools. Pair-level quality-control statistics were generated using pairtools stats and stored in a sorted pair file.

To remove duplicate contact pairs, rescued contacts were deduplicated based on contact coordinates and cell identifiers (R1, R2, R3, P1, and P2). Contacts mapped to standard chromosomes were retained. Contact records sharing the same composite identifier were considered duplicates, and a single representative contact was retained for each group. The number of reads contributing to each deduplicated contact was recorded as an additional count. The resulting unique contacts were balanced and stored in cool format using HiCExperiment and Cooler (v0.8.10) for visualization and downstream analysis. Cells with fewer than 10,000 unique contacts were excluded from further analyses.

### Single-cell A/B compartment analysis

Single-cell A/B compartment scores were calculated independently for each chromosome from ChromE-seq chromatin contact data at 100-kb resolution. The genome was partitioned into consecutive 100-kb bins using tileGenome. Because CpG density was subsequently used to orient compartment eigenvectors, bins containing no CpG dinucleotides were excluded. For each chromosome, intrachromosomal contacts were extracted and assigned to their corresponding 100-kb genomic bins. For each cell, a symmetric bin-by-bin contact matrix was constructed by including contacts in both orientations. A Pearson correlation matrix between genomic bins was then calculated. Undefined correlations were set to zero and diagonal values were set to zero. Bins without informative correlations, defined as having a row sum of zero, were excluded before eigendecomposition. To reduce the influence of extreme correlation values, correlation coefficients were winsorized at the 1st and 99th percentiles.

The first eigenvectors of the resulting correlation matrix were calculated using the symmetric eigensolver implemented in RSpectra. The eigenvector was normalized to unit length, scaled by the square root of the absolute value of its corresponding eigenvalue, and used as the single-cell compartment score. Because the sign of an eigenvector is arbitrary, compartment scores were oriented according to local CpG density, with positive values assigned to the CpG-rich, A-like compartment and negative values to the CpG-poor, B-like compartment. For cells in which eigendecomposition failed because of insufficient contact information, compartment scores were set to zero.

### Hi-C gene score

Chromatin contact-based gene scores were calculated using ArchR. Briefly, chromatin contact coordinates were used in place of ATAC-seq fragments as input to the ArchR gene score calculation. ArchR quantified gene-associated chromatin contacts using a distance-weighting scheme, which assigns greater weights to contacts proximal to gene bodies and transcription start sites, enabling the calculation of gene-level chromatin contact scores.

### Differential chromatin loop analysis

Differential chromatin looping between T21 and disomy cells was assessed using ChromE-seq contact counts. To maximize sensitivity and avoid restricting the analysis to loops identified in only one biological condition or cell population, candidate chromatin loops were called using the complete dataset as well as independently within each major cell type and genotype. Loop sets identified from these analyses were combined to generate a comprehensive union of candidate loops for downstream differential analysis. To account for uncertainty in loop-anchor positioning and local variation in contact enrichment, each identified loop anchor was expanded to include the immediately adjacent 10-kb genomic bins on either side. This generated nine bin-pair combinations within a 3 × 3 neighbourhood around each candidate loop. Each bin-pair combination was retained as a separate loop-associated feature rather than aggregating contact counts across the nine combinations. The resulting feature-by-cell contact-count matrix was imported into Seurat, normalized using NormalizeData with default parameters. Differential loop-associated contact frequencies between T21 and disomy HSC/MPP cells were assessed using FindMarkers function with a two-sided *t*-test. No minimum detection-frequency or fold-change thresholds were applied (min.pct = 0, logfc.threshold = 0), allowing all candidate bin-pair features represented in the matrix to be evaluated.

### ME-seq quality control, pre-processing, and visualization

Single-nucleus RNA-seq data generated by ChromE-seq and ME-seq were jointly processed using Seurat (v5). For quality control, ME-seq nuclei with fewer than 300 detected genes, more than 10,000 detected genes, or greater than 5% mitochondrial RNA content were excluded. ChromE-seq nuclei with fewer than 300 detected genes, fewer than 1,000 UMIs, or greater than 2% mitochondrial RNA content were excluded. Gene expression matrices from the two assays were combined and log-normalized using NormalizeData. The top 1,000 highly variable genes were identified using FindVariableFeatures, followed by data scaling with ScaleData and principal component analysis using RunPCA on the combined dataset. Batch effects across ChromE-seq and ME-seq libraries were corrected using Harmony. Cell clustering and UMAP visualization were performed using FindNeighbors and RunUMAP (dims = 1:35, min.dist = 0.3), respectively. To improve clustering and cell-type annotation, nuclei identified as doublets were removed before downstream analysis.

ME-ATAC-seq fragment files were processed using BPCells^25^. Cells with TSS enrichment scores greater than 4 and between 800 and 100,000 fragments were retained for downstream analysis. Gene scores were calculated using the gene_score_archr function.

Dimensionality reduction of chromatin accessibility and DNA methylation data was performed using cisTopic^26^. Open chromatin peaks were called using MACS2^27^ and standardized to 500-bp regions. For chromatin accessibility, the number of fragments overlapping each peak was quantified for each cell to generate a peak-by-cell accessibility matrix. For DNA methylation, the percentage of methylated cytosines within each peak was calculated for each cell to generate a peak-by-cell methylation matrix. These matrices were subsequently used for modality-specific dimensionality reduction and visualization. Topic models were trained independently on the accessibility and methylation matrices using runCGSModels (n_topics = 50) with default parameters, followed by UMAP embedding for visualization (min_dist = 0.2 for chromatin accessibility and min_dist = 0.3 for DNA methylation; n_neighbors = 100). In parallel, the percentage of methylated cytosines within gene bodies was calculated for each cell to generate a gene-by-cell methylation matrix for gene-level analyses.

### Transcription factor motif activity analysis

TF motif activity was quantified from single-cell ATAC-seq data using chromVAR. Briefly, TF binding motifs were identified in accessible chromatin regions using position weight matrices from the human transcription factor motif database. A peak-by-cell accessibility matrix was used to calculate chromVAR deviation scores, which quantify differences in chromatin accessibility at regions containing each TF motif relative to the expected accessibility. To account for technical biases, chromVAR generated background peak sets matched for GC content and average accessibility. Bias-corrected deviation Z-scores were calculated for each TF motif in individual cells and used to assess TF motif activity across cell types and experimental conditions.

### Label transfer from published haematopoietic references

To independently validate our manually assigned cell-type annotations, we compared our transcriptional profiles with published human haematopoietic reference atlases. We first mapped our cells to the human fetal liver haematopoietic reference generated by Popescu *et al.*^23^, comprising major haematopoietic stem, progenitor, and differentiated cell populations. Reference mapping and label transfer were performed using Seurat following the standard single-cell reference mapping workflow. Reference-derived cell-type assignments were compared with our manual annotations to assess concordance and refine ambiguous populations. As an additional independent reference, we performed label transfer using the BoneMarrowMap package^24^, following the authors’ recommended workflow and tutorial provided in the BoneMarrowMap GitHub repository. BoneMarrowMap-derived cell-type assignments were compared with our manual annotations using canonical marker-gene expression and confusion matrices to assess agreement between annotation strategies.

### Spatial transcriptomics quality control and pre-processing

Visium HD FASTQs and H&E whole-slide images were processed using 10x Genomics spaceranger (v4.0.1) to filter align sequencing data to the human genome (refdata-gex-GRCh38-2020-A), segment single cells based on H&E staining, and generate single-cell counts matrices by aggregating transcript counts from bins overlapping each segmented cell. To account for errors in H&E-based segmentation and transcript diffusion, ProSeg (v3.1.1)^28^, a transcript-based cell segmentation package, was run on spaceranger outputs. Counts matrices were filtered to keep cells with greater than 50 total counts, between 25 and 3,750 genes, less than 10% mitochondrial fraction, and less than 50% hemoglobin fraction. Genes expressed in less than 3 cells were removed. After filtering, data was preprocessed and integrated using rapids-singlecell (v0.15.0)^29^. Standard functions were used to log-normalize, identify highly variable genes, and scale data. Total counts, mitochondrial fraction, hemoglobin fraction, and cell cycle scores were regressed out, as this was found to improve noisy clustering due to red blood cell contamination and proliferation. Harmony integration through rapids-singlecell was performed^30^, using each individual sample as the batch, and clustering performed at resolution 0.8. Clusters were carefully annotated based on expression of specific marker genes.

### Spatial transcriptomics integration with single-cell data

We sought to improve data sparsity issues, and to reduce the computational resources required for integration, by first aggregating ChromE-seq RNA data into metacells using SEACells (v0.3.3)^31^, with a targeted 200 cells per metacell. Based on the assignment of cell barcodes into each RNA metacell, a Hi-C metacell matrix was also constructed through paired ChromE-seq data. The RNA metacells were projected onto Visium HD using TACCO (v0.4.0)^32^, and Hi-C scores from the same projected metacells were visualized, allowing for spatial inference of Hi-C signal on Visium HD data.

### Single-cell copy-number variation analysis

Infercnvpy (v0.6.1) was used to perform copy number analysis. For RNA data, transcripts were assigned back to genomic positions based on reference genome annotations. Counts data was log-normalized, then run through infercnvpy using a window size of 250, step of 10, and all disomic cells as reference. Hi-C data was processed similarly, by converting the 500 kb contact matrix into marginal contact frequency within each genomic bin followed by log-normalization. For Hi-C, infercnvpy was run using a window size of 25, step of 1, and all disomic cells as reference.

### GWAS trait enrichment analysis

Cell-type-specific enrichment of genetic variants associated with complex traits was assessed using gchromVAR^33^. For each cell type, chromatin accessibility profiles were aggregated across cells to generate pseudobulk profiles. GWAS summary statistics were used to assign trait-associated weights to accessible chromatin regions based on overlapping variants and their association statistics. gchromVAR was then applied to quantify enrichment of trait-associated variants within accessible regions for each cell type while correcting for technical and genomic biases, including overall accessibility, GC content and peak characteristics. Enrichment scores and corresponding significance values were used to identify cell types preferentially enriched for genetic risk associated with each trait.

### Pseudotime analysis and CytoTRACE 2

Developmental trajectories were reconstructed separately for the B-cell and erythroid lineages using Slingshot^34^ on the RNA-derived low-dimensional representation, with HSCs specified as the root population. Cells were ordered along each lineage trajectory and pseudotime was scaled from 0 to 100. RNA expression, chromatin accessibility, DNA methylation and Hi-C-derived regulatory features were subsequently summarized along pseudotime to compare the temporal ordering of regulatory changes during differentiation.

Cellular differentiation potential was independently estimated from the RNA expression matrix using CytoTRACE 2^18^. Potency scores and predicted potency states were compared across haematopoietic cell types and between Disomy and T21 cells. Higher CytoTRACE 2 scores indicate greater developmental potential, whereas lower scores indicate more differentiated cellular states.

### Peak–gene *cis*-regulatory association analysis

ATAC-defined accessible peaks were used as the common set of candidate *cis*-regulatory elements. The peaks are called using MACS2 and normalized to 500 bp width. Candidate peak–gene pairs were generated by pairing peaks with protein-coding genes within 2 Mb of transcription starting sites. Peak–gene associations were identified by Spearman correlation between regulatory-element activity and gene expression across matched single-cell profiles. Significance was evaluated relative to background regulatory elements matched for features including GC content, average chromatin signal and gene-expression level. ATAC–RNA associations were evaluated for positive regulatory relationships, whereas mCG–RNA associations were evaluated using a two-sided test to capture both positive and negative relationships between DNA methylation and transcription. Analyses were performed across the entire dataset to identify shared and lineage-restricted regulatory interactions. Genes associated with multiple significant regulatory peaks within the proximal *cis*-regulatory window were classified as domains of regulatory chromatin (DORCs)^3^ and used for subsequent regulatory-network analysis.

### Loop-assisted DORC analysis

Loop-assisted domains of regulatory chromatin (LA-DORCs) were identified by integrating chromatin accessibility, DNA methylation, gene expression and chromatin-contact information. Candidate *cis*-regulatory elements were first linked to nearby genes using cell type-specific peak–gene associations derived independently from ATAC–RNA and mCG–RNA correlations. Significant regulatory element–gene pairs supported by both chromatin modalities were then intersected with chromatin loops detected by ChromE-seq, retaining associations in which the regulatory element and target gene promoter were connected by a chromatin contact. Genes supported by multiple loop-linked regulatory elements were defined as LA-DORCs. LA-DORC scores were calculated by aggregating the corresponding regulatory signals for each gene and were used to characterize lineage-specific regulatory programmes and construct gene regulatory networks.

### Gene regulatory network analysis

TF regulatory networks were reconstructed by integrating TF expression, TF motif information and chromatin states at DORC-associated regulatory elements. TF motif occurrences were identified within ATAC-defined regulatory peaks using the human motif collection from chromVARmotifs and motifmatchr. ATAC-based TF motif activity was estimated using chromVAR with GC- and accessibility-matched background peaks. Methylation-associated TF motif activity was calculated over the same peak coordinates using an adapted chromVAR framework in which local cytosine coverage and regional methylation levels were incorporated into the expected motif signal.

TF–DORC interactions were inferred using FigR^35^. For each DORC, FigR integrates enrichment of a TF motif within its linked regulatory peaks with the correlation between TF expression and DORC regulatory activity across cells. ATAC DORC activity was calculated from accessibility across linked peaks (*FDR*<0.05), whereas mCG DORC activity was calculated from methylation levels across the same regulatory elements. The resulting signed regulation score represents both the strength and direction of the inferred TF–target relationship, allowing regulatory interactions to be classified as ATAC-associated activation or repression and mCG-associated activation or repression. ATAC- and mCG-derived interactions were combined to generate the multimodal chromatin–mCG–RNA regulatory network.

To determine how trisomy 21 alters regulatory circuitry, networks were reconstructed independently in Disomy and T21 cells using the same TF–DORC framework. For each TF, regulation scores in the two conditions were compared across its target DORCs, and network perturbation was quantified by deviation from the Disomy–T21 diagonal. TFs showing systematic changes in regulation scores were prioritized as candidate T21-disrupted regulators. DORC targets overlapping genes associated with T21-perturbed chromatin contacts were additionally annotated in the network to connect changes in TF regulation with alterations in 3D genome organization.

### CRISPR/Cas9-induced chromatin loop deletions

Guide RNAs targeting the T21-sensitive and non-differential *EBF1* loop anchors were designed using Benchling. For each anchor, candidate guide RNAs were selected on both the upstream and downstream sides of the targeted region, within approximately 250 bp of the anchor. Candidate guides per anchor were shortlisted using Benchling on-target scores >45^36^ and off-target scores >40^37^ and were experimentally evaluated in human fetal liver-derived CD34^+^ HSPCs following electroporation by PCR amplification of the targeted region, Sanger sequencing (GENEWIZ) and ICE analysis^21^. The highest-efficiency upstream and downstream guides for each anchor were selected for subsequent experiments. Two guide RNAs targeting the olfactory receptor gene *OR2W5* were used as a negative control^38^ (**Extended Data Table 10)**.

CRISPR/Cas9 ribonucleoprotein (RNP) complexes were assembled using chemically synthesized Alt-R CRISPR/Cas9 crRNAs and tracrRNA (Integrated DNA Technologies, IDT) together with recombinant Cas9 nuclease (IDT). crRNAs and tracrRNA were reconstituted to 200 μM in TE buffer (IDT). For each target locus, the upstream and downstream crRNAs were combined with tracrRNA at a molar ratio of 0.5:0.5:1 and annealed at 95 °C for 5 min, followed by cooling to room temperature. The two guides were used simultaneously to generate a deletion spanning the targeted loop-anchor region.

For each electroporation reaction, 1.2 μL of annealed crRNA–tracrRNA complex, 1.7 μL Cas9 protein and 2.1 μL PBS (Cytiva) were combined in a low-binding microcentrifuge tube and incubated at room temperature for 15 min to assemble the RNP complex. Electroporation enhancer (1 μL of 100 μM stock; IDT) was then added. Precultured cells were washed with prewarmed PBS and centrifuged at 350 × *g* for 10 min. Approximately 2.5 × 10^5^ cells were resuspended in 20 μL P3 Primary Cell Nucleofector Solution (Lonza) per reaction, combined with the RNP complex and transferred to a 4D-Nucleofector cuvette. Cells were electroporated using a 4D-Nucleofector (Lonza) with programme DZ-100.

For xenotransplantation experiments, human FL CD45^+^CD34^+^ HSPCs were isolated by fluorescence-activated cell sorting on day 1 and allowed to recover in culture in serum-free X-VIVO 10 medium (Lonza) as previously described^38^. Cells were electroporated with CRISPR/Cas9 RNP complexes on day 3 and transplanted into NSG mice on day 4. An aliquot of the cells used for transplantation was retained for genotyping. Editing efficiency at the targeted loci was assessed by PCR amplification of the edited region, Sanger sequencing (GENEWIZ) and ICE analysis.

### Xenotransplantation and flow cytometry analysis

Following electroporation, cells were allowed to recover overnight at 37 °C in a humidified tissue-culture incubator. The following day, 1 × 10^5^ edited cells were transplanted by intrafemoral injection into 8- to 12-week-old female *NOD.Cg-Prkdc^scid^Il2rg^tm1Wjl^/SzJ* (NSG) mice (The Jackson Laboratory) that had received sublethal irradiation (230 cGy) 24 h before transplantation. Littermates were randomly assigned to experimental groups.

Mice were euthanized 12 weeks after transplantation. Bone marrow was collected from the left and right femurs and tibias and flushed with a staining buffer (PBS containing 2.5% FBS). Cells were centrifuged at 350 × *g* for 10 min and resuspended in a 500 μL staining buffer. Human engraftment was assessed by flow cytometry and was considered successful when human CD45^+^ cells represented >2% of live singlets. Only mice meeting this prespecified engraftment criterion were included in downstream analyses.

For flow-cytometric analysis, 50 μL of cell suspension was combined with 50 μL of antibody mixture and incubated for 60 min at 4 °C protected from light. Cells were washed once with a staining buffer and resuspended in a 250 μL staining buffer before acquisition on a BD LSRFortessa (BD Biosciences). A pan-lineage panel comprising CD45, CD19, CD3, CD33, CD41, CD235a, CD71, CD34 and CD117 was used to quantify human engraftment and major haematopoietic lineages. A lymphoid-progenitor panel comprising CD45, CD34, CD38, CD45RA, CD10, CD19, CD33, CD235a and CD20 was used to resolve early lymphoid and B-cell developmental populations. Antibody clones, fluorochromes, dilutions and suppliers are provided in **Extended Data Table 9**. The same staining panels were used for analysis of primary fetal liver samples. Remaining unstained cells from each xenograft were cryopreserved and stored in liquid nitrogen.

Viable singlets were identified before lineage gating. Human engraftment was defined by CD45 expression. Early lymphoid populations were quantified within the CD33^−^CD235a^−^CD45^+^CD34^+^ compartment using CD38, CD45RA, CD10 and CD19 expression (**Extended Data Fig. 7a**). MLPs were defined as CD38^−^CD45RA^+^ cells. CLPs were defined as CD38^+^CD19^−^CD10^+^ cells, whereas committed B-cell progenitors were defined as CD38^+^CD19^+^CD10^+/−^ cells. The B-cell progenitor gate comprises the combined pre-pro-B/pro-B population used throughout this study.

### Immunohistochemistry staining and quantification

CD19 immunohistochemistry was performed on 4-μm sections obtained from the same FFPE fetal liver blocks used for Visium HD profiling. Staining was performed using the Leica Bond RX automated staining platform (Leica Microsystems) according to a standardized automated workflow. Sections were deparaffinized and subjected to heat-induced epitope retrieval. Endogenous peroxidase activity was blocked before incubation with an anti-CD19 primary antibody (Agilent Technologies, LE-CD19). Bound antibody was detected using a polymer-based horseradish peroxidase detection system with 3,3′-diaminobenzidine (DAB) as the chromogen, followed by haematoxylin counterstaining. Slides were dehydrated, coverslipped and scanned at 40× magnification.

CD19-stained fetal liver sections were analysed in ImageJ. For each liver, five representative regions of interest were selected for quantification, with more than 1,100 total cells counted per specimen. Color deconvolution was performed using the Color Deconvolution 2 plugin to separate haematoxylin and DAB signals. Nuclei were segmented from the haematoxylin channel, and CD19-positive cells were identified based on DAB staining. The percentage of CD19^+^ cells was calculated as the number of CD19^+^ cells divided by the total number of segmented cells across the analysed ROIs for each specimen.

## Data availability

Raw and processed sequencing data will be released publicly upon publication.

## Code availability

We have assembled a web-based Shiny interface to interact with the data in a user-friendly manner for the public, which can be accessed here: https://malabsinai-nas.synology.me/fetal_liver/.

Additionally, we have hosted the ATAC tracks (pseudobulked per celltype) in a compatible format for the UCSC Genome Browser, which can be accessed here: https://genome.ucsc.edu/cgi-bin/hgTracks?db=hg38&position=chr5:157500000-159500000&hubUrl=https://malabsinai-nas.synology.me:8443/hub.txt.

Code for aligning ME-seq data can be found here: https://github.com/sai-ma-group/ME-seq-alignment-pipeline. Code for aligning ChromE-seq data is currently in preparation, and will be available on publication of this manuscript.

## Acknowledgements

We thank all the members of the Ma and Wagenblast labs for critical reading of the manuscript and helpful discussions. All research involving human fetal tissue from elective abortions was completed before 22 January 2026, the effective date of NIH Guide Notice NOT-OD-26-028. No NIH funds were used for research involving such tissue after that date, and subsequent work was limited to analysis of existing data and preparation of this manuscript. S.M. acknowledges support from the National Institutes of Health (NIH) through awards U01CA290442 and R61CA297881. S.M. and E.W. acknowledge support from the NIH R21CA301237 and HMTF Pilot Award Program for MDS Research. E.W. acknowledges support from the NIH through award R01CA290681 and support through a Blood Cancer United (formerly Leukemia and Lymphoma Society) Specialized Center of Research Program Grant (7039-25). In addition, E.W. is a Pew-Stewart Scholar supported by the Pew-Stewart Scholars Program for Cancer Research. M.Z. is supported by the American Society of Hematology (ASH) Graduate Hematology Award. B.Y.S. is supported by the NIH Medical Scientist Training (T32GM146636) and a National Cancer Institute predoctoral fellowship (F30CA301790). This work was supported in part by the Bioinformatics for Next Generation Sequencing (BiNGS) shared resource facility within the Tisch Cancer Center at the Icahn School of Medicine at Mount Sinai, which is partially supported by NIH grant P30CA196521, and the Icahn School of Medicine at Mount Sinai (ISMMS) Skin Biology and Disease Resource-based Center NIAMS P30 (AR079200). This work was also supported in part through the computational resources and staff expertise provided by Scientific Computing at the Icahn School of Medicine at Mount Sinai supported by the Clinical and Translational Science Awards (CTSA) grant UL1TR004419 from the National Center for Advancing Translational Sciences. Research reported in this publication was also supported by the Office of Research Infrastructure of the National Institutes of Health under award number S10OD038231.

## Additional information

### Contributions

Z.L. and S.M. developed ChromE-seq protocol. Z.L. generated the single-cell sequencing library. M.Z. performed sorting and sample processing for the fetal liver samples and performed functional experiments. K.W., M.Q.A. and P.F. assisted with xenotransplantation experiments. Z.L., B.Y.S., S.M., L.L.B., I.O., Z.W, D.H performed computational analysis. B.Z. assisted with ME-seq library preparation. Z.L., M.Z., B.Y.S participated in writing the manuscript. E.W. and S.M. conceived and supervised the research, and all authors reviewed the manuscript.

